# The value structure of metabolic states

**DOI:** 10.1101/483891

**Authors:** Wolfram Liebermeister

## Abstract

To improve their metabolic performance, cells need to find compromises between high metabolic fluxes, low enzyme investments, and well-adapted metabolite concentrations. In mathematical models, such compromises can be described by optimality problems that trade metabolic benefit against enzyme cost. While many such modelling frameworks exist, they are often hard to compare and combine. To unify these modelling approaches, I propose a theory that characterises metabolic systems by a value structure, that is, a pattern of local costs and benefits assigned to all elements in the network. The economic values of metabolites, fluxes, and enzymes are interlinked by local balance equations. Formally defined as shadow values, the economic variables serve as local proxies for benefits that arise anywhere in the network, but are represented as local costs or benefits in the reaction of interest. Here I derive economic variables and their balance equations for kinetic, stoichiometric, and cell models. Metabolic value theory provides a new perspective on biochemical networks, defines concepts for comparing and combining metabolic optimality problems, and is useful for semi-automatic, layered, and modular modelling.

## 1 Introduction

A given metabolic network can carry various combinations of metabolic fluxes, metabolite concentrations, and enzyme levels [1]. Which pathway fluxes should a cell use in a given moment, and why, to obtain a selection advantage? How should it allocate its protein budget to metabolic reactions and how should protein levels be adapted when cells are perturbed [2]? Should an inhibited enzyme be overexpressed (to compensate for the inhibition) or should the entire pathway be switched off (because now it works inefficiently)? And depending on what should this decision be taken? If we ask such questions, we do not only ask how cells behave in reality, but how they *should* behave in order to maximise their selection advantage. To answer this, studying mechanisms – how enzyme activities are regulated biochemically – is not enough: we need to consider enzyme function: e.g. how particular pattern of enzyme levels and fluxes contribute to fitness. Mathematically, the functioning of metabolism can be understood through economic principles, describing how cells may maximise metabolic performance. This leads to notions of cost and benefit: proteomics data (Fig. 1) can be seen as the cell’s investome, where proteins that are highly expressed – and therefore costly – are expected to provide a high benefit. To define optimal enzyme levels in a given kinetic model (leading to optimal fluxes and metabolite concentrations) [3, 2, 4, 5], enzyme profiles may be scored by a benefit function – usually a function of steady-state fluxes and metabolite concentrations – and be penalised or constrained by a cost function [3, 4], for example the total enzyme concentration [6]. In such models, aside from network structure (the “blueprint”) and metabolic dynamics (the “kinetics” of of compound concentrations and reaction rates), we introduce a third layer of analysis, a layer concerning biological function, physiology, or as I call it, “metabolic economics”. Of course, all three layers – structure, dynamics, and economics – are tightly related.

**Figure 1.**
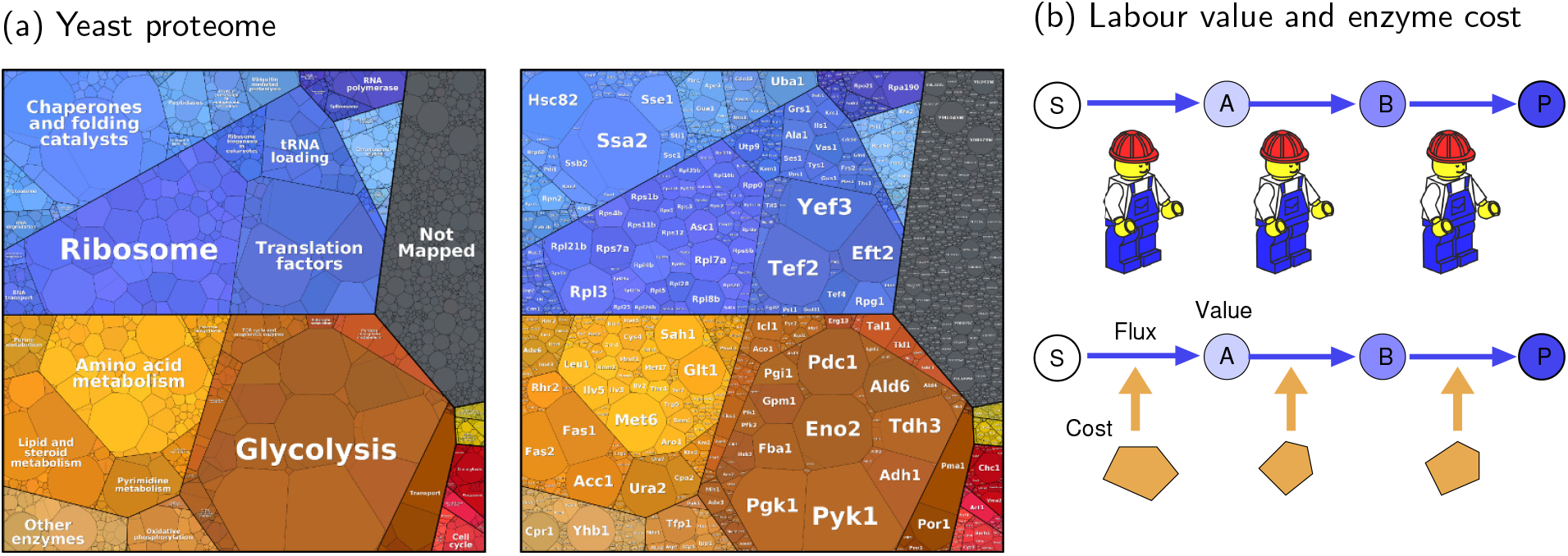
Protein investments in cells. (a) Proteome of the yeast *S. cerevisiae*. Protein abundances are shown as polygon areas (proteomap from [11], original data from [12]). A large fraction of the protein budget is spent on metabolic enzymes (yellow, orange and brown). To explain the abundance of individual enzymes, we associate proteome mass fractions with costs and assume that these costs must be justified by equally large benefits (describing the enzymes’ contributions to cell fitness). Here I show how notions of “cost” and “benefit” can be precisely defined, assuming that enzyme allocation to reactions and pathways should maximise fitness. (b) In economics, labour value theories define the value of a commodity (shades of blue) by the labour time invested in its production (plus investments in material and machines). Along a production chain, the value increases in each step because of the labour invested. I propose a similar picture for metabolism: each metabolite has an embodied investment, representing the total enzyme labour (and precursor value) invested in its production. This means: to justify its own cost, each enzyme must produce more valuable metabolites from less valuable ones, and the value difference in a reaction (including cofactors), multiplied by the flux, must be equal to the enzyme investment in this reaction. The values increase along a pathway corresponding to accumulating enzyme investments.

In economic cell models, we assume that cells have certain tasks and that their performance can be quantified by a utiliy function called “fitness”. This “metabolic objective” – for example, biomass production per cell volume – requires well-adjusted fluxes, metabolite concentrations, and enzyme levels in the entire network, which gives reactions, metabolites, and enzymes a “use value”. The value structure – the pattern of all metabolite, flux, and enzyme values across the network – reflects the way the physical cell variables (concentrations and fluxes) interact or constrain each other. Large metabolic fluxes support biomass production but also require high substrate and enzyme levels: the enzyme molecules occupy valuable space [7] and their production and maintenance consume resources such as amino acids, ribosomes, and energy. All these requirements make fluxes costly. Conversely, if glucose import contributes (indirectly) to ATP production, this makes the transporter (indirectly) beneficial. In bacteria, protein costs and benefits have been quantified by measuring cell growth after a forced expression of protein [4, 8]). To maximise metabolic performance, cells must prioritise the usage of metabolic pathways and adjust their fluxes to changing supplies and demands. The resulting fluxes reflect a compromise between tasks – e.g. the production of biomass components – and a protein cost or limited protein capacity. By regulating their enzyme levels, cells can save protein cost, disrupt futile cycles that waste valuable metabolites [9], and adapt their fluxes to different objectives, for example, maximal cell growth or product yield [10, 5].

To model all this, we may imagine a kinetic model of the entire cell, describing resource allocation in metabolism and macromolecule production. However, with realistic kinetic rate laws such models would lead to large, non-convex optimality problems that are hard or impossible to solve. This is why existing modelling frameworks use simplifications: some focus on smaller pathways (with enzyme levels as control variables, bounded or penalised by costs, while others (like RBA or FBA with molecular crowding) replace enzyme kinetics by “capacity constraints” based on fixed enzyme efficiencies, or ignore it completely (such as classical FBA). However, all these models should portray the same reality! If we assume that an enzyme is “costly”, this cost should be comparable between models: cost should not only be a variable within a model, but should be a real property of the biological system described, i.e. comparable between models. To see how different models can be compared and combined, we need a general theory and unified formulae.

In [13], I described optimality problems with different types of state variables (fluxes, metabolite concentrations, and enzyme levels) and physical and biochemical laws (rate laws, stationarity, and physiological bounds). Metabolism contributes to cell fitness, and we can quantify this contribution by benefit and cost terms (flux benefit, metabolite cost, and enzyme cost). Obviously, benefits should be high and costs should be low, and in metabolic models we can describe this by optimising them or constraining them by lower and upper bounds. Cost/benefit compromises can be modelled in different ways: metabolic targets can be combined to a single objective, or one target is optimised while the others are fixed or constrained. In this way, we can construct different optimality or satisfiability problems and obtain a variety of models (including the various forms of FBA) that can be optimised numerically. However here, instead of solving such problems one by one, we may also ask more general questions. Are there patterns of fluxes or enzyme investments that cannot be optimal in any possible model? If we observe an enzyme investome (e.g. proteomics data interpreted as costs), can we tell if it represents an optimal state? How do optimal enzyme investments reflect metabolic network structure, enzyme kinetics, and enzyme prices (e.g. their molecular weights)? To answer such questions, we need a theory of metabolic value. The theory should hold for simple pathways with a single product, as well as for the entire metabolic network, with branched and cyclic fluxes and biomass precursors as the output. To obtain such a theory, we start from existing optimality problems, consider their optimality conditions, and describe them by general formulae. As a result, we obtain general, local laws for optimal metabolic states.

Metabolic economics is a theory of metabolic optimality problems as described in [13]. As an optimality condition, all these models imply a balance between cost and benefit gradients. Here I describe the same balances as a “value structure”, a pattern of costs and benefits of individual network elements, and propose a theory for this structure. Metabolic value theory uses *economic* variables to describe the importance of enzymes, reactions, and metabolites for a cell. Mathematically, these economic values are dual variables that arise from the optimality problems as Lagrange multipliers or shadow prices. In practice, they characterise the economic value of fluxes, enzymes, and metabolites for a cell, in the context of a given optimality problem. Using these values, optimality conditions can be written as balances between local values: for example, between the values of an enzyme and its catalysed reaction, between the value of a reaction and of its substrates and products, or between the values of a metabolite and the reactions it influences kinetically (as a reactant or allosteric regulator). In the network, these balances generate what I call a “value structure”: a pattern of values assigned to reactions, metabolites and enzymes in the entire network, and interdependent between neighbouring network elements. These dependencies can also be described by a “flow of value”. For example, a balance equation states that in each reaction the incoming value (substrate or enzyme investments) must match the outgoing value (production of valuable products). Due to the inflow of enzyme investments in every reaction, the value of metabolites (shades of blue) increases along the flux. A production chain as shown in Figure 1 (b) can be a simple metabolic pathway, but may also represent the entire cell, with three reactions subsuming all transporters, all metabolic reactions, and all steps of protein synthesis. Economic variables and balance equations arise in the context of different optimality problems and provide a general framework for optimal metabolic states.

In this paper, I outline metabolic value theory for metablic optimality problems, following the concepts and terminology from [13]. In section 2 I introduce economic variables and their laws as a postulate. In section 3, I derive these variables and laws from existing optimality problems. In section 4, I describe this as a general theory. Details on optimality problems, derivations of economic laws, and mathematical proofs can be found in the SI.

## 2 Metabolic value theory

### 2.1 Metabolic optimality problems and their optimality conditions

Many metabolic optimality problems follow from a simple general problem: maximising a fitness function on the set of feasible metablic states [13]. Kinetic models are described in Box 1: metabolic states are described by fluxes **v**, metabolite concentrations **c**, and enzyme levels^1^ **e**. They need to respect stationarity, enzymatic rate laws, and physiological bounds, and maybe a predefined flux benefit or other extra constraints. The feasible states (**v, c, e**) form an algebraic manifold called state manifold. To model trade-offs between metabolic targets (e.g. between flux benefit, metabolite cost, and enzyme cost), we assume a fitness function

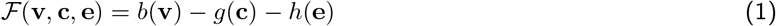

composed of benefit and cost terms for the vectors **v** (fluxes), **c** (metabolite concentrations), and **e** (enzyme levels). This fitness needs to be maximised while our variables **v, c**, and **e** are constrained by physical and physiological laws (mass balance, enzymatic rate laws, dilution effects, and density constraints). Other optimality problems set objectives and constraints differently: for example, we may minimise cost at a given benefit, maximise benefit at a given cost, use multi-objective optimisation, and may consider additional constraints or objectives. However, all these problems – single-objective, constrained, and multi-objective problems – lead to optimality conditions of the same form [13]: the gradients of metabolic targets in the space of control variables must be balanced, that is, a convex combination of these gradients must vanish. Unfortunately, some gradients are hard to compute: for instance, to obtain a flux benefit gradient in the space of enzyme levels, we need to express the fluxes as functions of enzyme levels, which cannot be done separately for each reaction, but requires knowledge about the entire network. This is no surprise: to understand how a changing enzyme level changes the metabolic benefit, we need to trace its network-wide effects.

#### Box 1

Metabolic optimality problems

##### Kinetic metabolic model in standard form: physical variables and their dependencies

In metabolic models, the elements of a biological network (metabolites, reactions, enzymes, and others) are associated with physical variables such as concentrations, fluxes, or production rates. Kinetic metabolic models in a “standard form” contain foru main types of variables, shown below. The variables are dependent and subject to constraints: concentrations determine reaction rates, and reaction rates determine metabolite net production which can either change metabolite concentrations in time (to model dynamic behaviour) or are set to zero by a stationarity constraint (to model steady states).

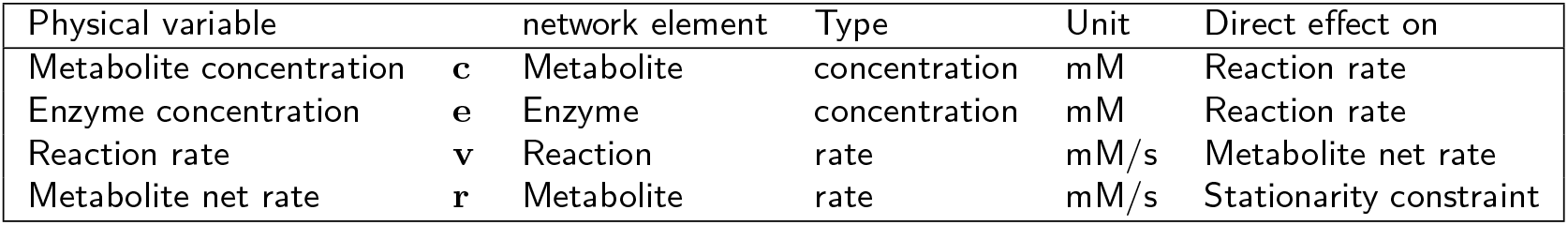

##### Metabolic optimality problems: bounds, constraints, and objectives

Figure (a) shows the physical variables in kinetic metabolic models (metabolite and enzyme levels; reaction rates; and metabolite production rates). For convenience, net production rates of metabolites are treated as separate variables.

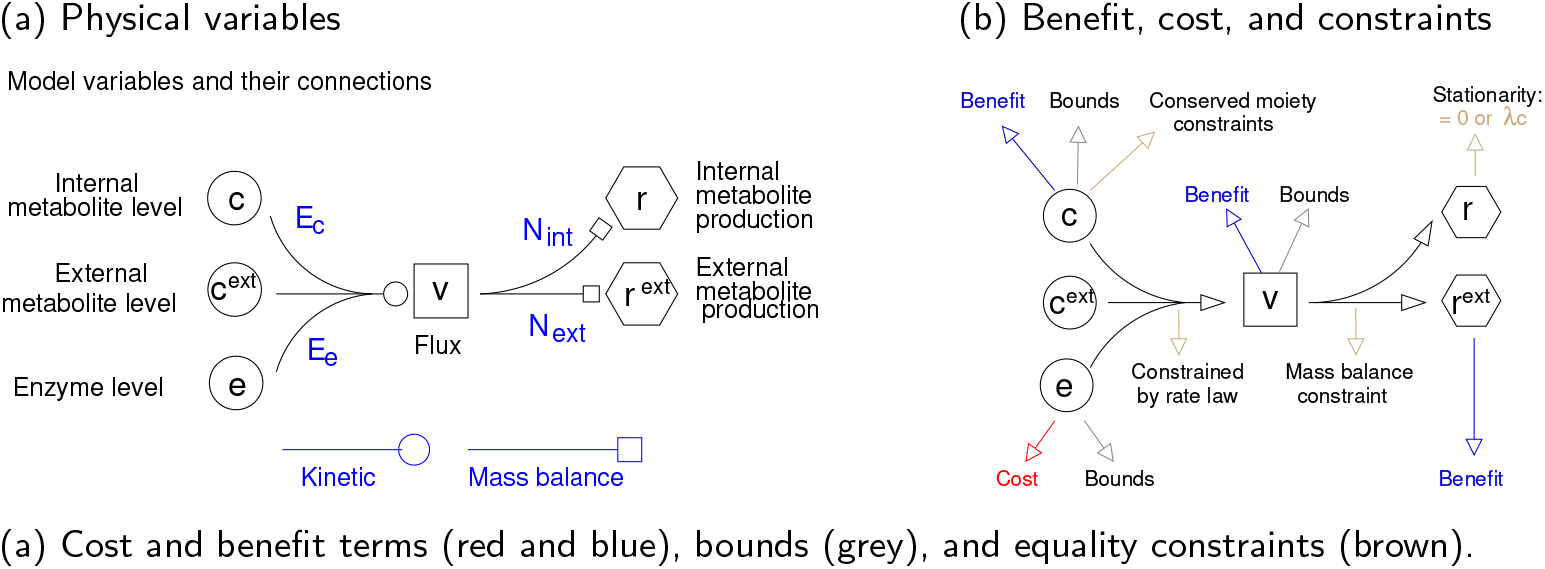

In the “standard form” of such models, we treat enzyme and metabolite concentrations as free variables, which determine the fluxes. Arrows represent direct dependencies, given by kinetic rate laws **v**(**c, e**) and mass balances **r**(**v**) = **N v**. The derivatives of these functions form the elasticity matrices **E**_c_ and **E**_e_ and the stoichiometric matrix **N**. These connection matrices can be used to propagate variations, e.g. *δ***v** = **E**_e_ *δ***e** + **E**_c_ *δ***c**. To couple different compound concentrationsby constraints, we may describe weighted sums or differences of compound concentrations by auxiliary variables (e.g. “total mass density”).

Since changes of enzyme levels change the steady-state fluxes “non-locally” in the entire network, it is no surprise that also the economic variables, which score these actions by fitness effects, follow “non-local” laws. Starting from a given optimality problems (e.g. with metabolite or enzyme levels as control variables), the optimality condition leads economic laws that link physical variables (concentrations, fluxes, enzyme levels) to their dual economic variables (value and price). For example, the balance of gradients (in enzyme space) of flux benefit, (negative) metabolite cost, and (negative) enzyme cost, can be written as a balance equation^2^

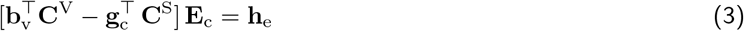

relating “enzyme prices” (elements of the vector **h**_e_ on the right) to “enzyme use values” (the elements of the vector given by the formula on the left). The symbols in the formula denote enzyme elasticity coefficients (matrix **E**_c_ = Dg(*v*_*l*_*/e*_*l*_)), control coefficients (matrices **C**^V^ and **C**^S^), and fitness derivatives (flux gains in **b**_v_ = *∂*𝓕*/∂***v**, metabolite prices in **g**_c_ = −*∂*𝓕*/∂***c**, enzyme prices in **h**_e_ = −*∂*𝓕*/***e**). But a problem remains. The equations couple the variables non-locally: the control coefficient matrices describe how a variable (e.g. a flux *v*_*l*_) affects distant variables in the network in steady state, (e.g. a concentration *c*_*i*_, associated with a concentration price 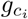). This is no surprise, because the need for enzyme adjustments in a reaction depends on a need for flux or concentration changes elsewhere. However, these network-wide dependencies are hard to understand and they even pose a problem for models in general: if the economic values in a pathway depend on cost and benefit elsewhere, how can we model this pathway without describing the entire cell? Can we optimise a pathway in isolation? While this may sound obvious – because modellers do this all the time – the question behind it is: are our pathway models based on ad-hoc assumptions, or can they be rigorously derived from a larger theory, in which fitness objectives (for pathways) represent what they should, namely contributions to cell fitness.

Here I propose a solution: all fitness effects in the rest of the cell, outside the pathway, are captured by proxy variables on the pathway boundary! By defining such variables and to show that they can capture all network-wide fitness effects, we obtain a local theory – a theory in which network-wide fitness effects are represented by local economic values. How does this work? In Eq. (3), the bracket term on the left can be rewritten as a simple sum 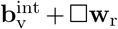, where 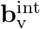 describing a fitness effects and □**w**_r_ is a difference of local variables called economic potentials, which describe the “use values” of individual reactants. The resulting balance equation

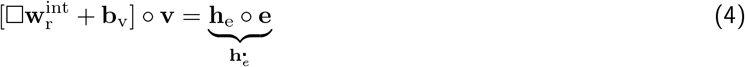

does not contain control coefficients anymore. The equation contains only local variables referring to a reaction, its enzyme, and its reactants. Below I show how such balances (and a theory of such balances) can be derived. I start from metabolic optimality problems [13] and rewrite them in “expanded form”, in which all dependencies between variables appear as explicit constraints, connecting neighbouring network elements. The constraints are associated with shadow values, which we see as economic variables, and the optimality conditions – which relate shadow values locally in the network – yield the basic laws of our theory. Metabolic value theory holds for a wide range of model types and applies to metabolic pathways, metabolic networks, and systems beyond metabolism.

### 2.2 Economic values and local economic rules

Before going into details, let me introduce some main concepts of metabolic value theory. The “economic side” of metabolic states is governed by economic values and general economic laws. In kinetic models, we consider economic potentials 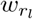, metabolite loads 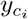, and enzyme prices 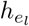. The economic variables represent economic values associated with the physical variables – for example, 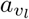 is a flux burden associated with flux *v*_*l*_ and thus with reaction *l*. While flux gains and metabolite prices (inclusing shadow values from flux and metabolite bounds) describe “direct” effects of the fitness, potentials and loads describe indirect effects, mediated through the metabolic steady state. The symbol Δ denotes the difference between products and substrates of a reaction, defined as Δ*x*_*l*_ = ∑_*i*_ *n*_*il*_ *x*_*i*_, with the stoichiometric coefficients *n*_*il*_. The (unscaled) elasticities 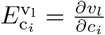 depend on the rate laws and on the metablic state.

The metabolic values obey basic laws called economic rules. An economic rule relates an economic value (of a reaction, metabolite, or enzyme) to the economic values around it. To see how these rules emerge, we can study small perturbations of a given metabolic state and assess their fitness effects. In metabolic models, variables are coupled through physical laws (e.g. a reaction rate depends on metabolite and enzyme concentrations, and a metabolite net rate depends on the rate of producing and consuming reactions. Accordingly, variations in a variable influence neighbour variables via “connections” such as elasticity cofficients or stoichiometric coefficients. For example, starting from a given steady state (**v, c, e**), a perturbation of metabolite concentrations *δ***c** and enzyme levels *δ***e** will lead to (immediate) changes in reaction rates, metabolite net rates, and fitness function:

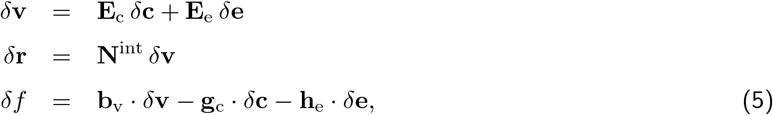

with the elasticity matrices **E**_c_ and **E**_e_ and the stoichiometric matrix **N**^int^ as connection matrices, and with the gain and price vectors **b**_v_, **g**_c_, and **h**_e_. However, we are not interested in local, immediate changes, but in long-term, network-wide changes of steady states, caused by indirect effects in the entire network. In such steady state changes, *δ***v**, *δ***c**, and *δ***e** are dependent. For example, a perturbation *δ***c**^pert^ and *δ***e**^pert^ for the concentrations will lead to long-term steady state changes *δ***c**^st^ and *δ***v**^st^, satisfying **N**^int^ *δ***v**^st^ (stationarity) and **G** *δ***c**^st^ = **G** *δ***c**^pert^ (moiety conservation). Altogether, our fitness change reads

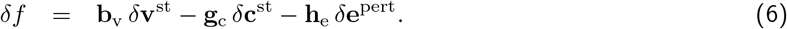

To compute it, we already need to know the steady-state changes. In contrast, we can also write the fitness change directly in terms of our original perturbations. For generality, we set *δ***v** = **E**_c_ *δ***c**^pert^ + **E**_e_ *δ***e**^pert^ + *δ***v**^pert^.

Written in terms of our original perturbation, the same fitness change reads

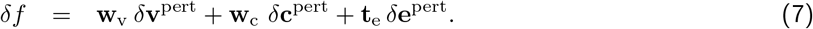

How can the “economic values” in the vectors **w**_v_, **w**_c_, and **t**_e_ be determined? While this may be difficult, we will see that these values – even if they may be unknown – are subject to general laws, called economic rules, that interconnect economic values between neighbouring enzyme levels, metabolite concentrations and flux in the metabolic network. To obtain these rules, we first note that each physical variable *x* can contribute to fitness in two ways. If *x* appears in the fitness function, it carries a direct value^3^ *f*_**x**_ = *∂f/∂x*. But it may also influence some neighbour variables; if these variables have a (direct and indirect) effect on fitness, this gives *x* an indirect value, which depends on the neighbour variables’ value and on *x*’s influence on that neighbour variable. Direct and indirect value together yield the total value of *x*. The economic rules state this mathematically. Let us see an example. The phosphofructokinase reaction in glycolysis, F6P + ATP ⇔ FBP + ADP, appears in the mass balances of F6P, ATP, FBP, and ADP. If these mass balances are described by explicit constraints, the corresponding shadow prices can be seen as economic potentials 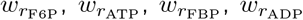. The economic value 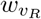 of our flux (associated with the flux *v*_*R*_), represents fitness effects of this reaction that are caused by the production and consumotion of these metabolites. In addition, the value *w*_v_ may contain an effective direct flux gain abd a shadow value caused by a flux bound (the latter two terms can be seen as an effective direct flux gain). Altogether, we obtain five terms, which can be grouped into indirect and direct values:

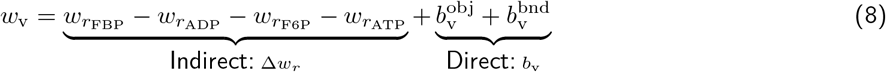

In ths formula, the indirect economic value of our reaction flux is acquired from the production values of the metabolites. The prefactors (+1 and -1), given by stoichiometric coefficients, are the same numbers that also connect variations of the physical variables (flux and production rates).

#### Economic rules: metabolic value consist of direct and acquired (indirect) values

Each dynamic variable is associated with an economic value, and each economic value consists of a direct and an indirect part. The direct value is acquired directly from the fitness function, while the indirect part is acquired from neighbouring (or more precisely: child) variables. Formally, this is expressed by the economic rules. They read, in vectorial form

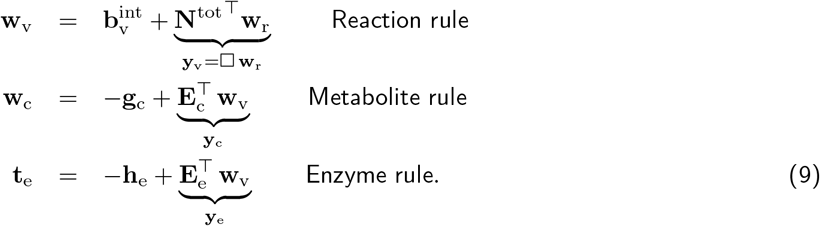

In each rule, the total value (on the left) is equated to a sum of direct and indirect values (the two terms on the right). Direct values (or “gains”) arise from direct effects on the fitness and from shadow values for bounds. Indirect effects (between a variable and the benefit function) arise from our variables acting on neighbouring variables and from these neighbouring variables’ indirect action on benefit. This is why also the indirect values (or “loads”) are acquired from these elements: they can be computed from the connections to these elements and from their economic values.

All three rules have the same form, splitting a total value into direct value and an indirect value. The *direct values* (flux gains **b**_v_, metabolite prices **b**_c_, and enzyme prices **h**_e_) are partial derivatives of the fitness function and exist for any physical variables with direct effects on fitness. The indirect values (economic potentials 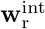, flux demands **w**_v_, and concentration demands **w**_c_) arise from dependence constraints caused by namely stationarity, rate laws, and moiety conservation. For example, a flux gain 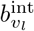 in a reaction contributes to the flux value 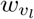. Each rule links an economic value (flux value 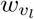, metabolite value 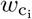, or enzyme price 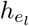) to the corresponding direct value and to economic variables in child variables (in adjacent network elements). The flux values 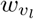 are already knwon from above. Concentration values are defined as a vector **w**_c_ = **G**^*T*^ **y**_cm_, where **y**_cm_ contains the Lagrange multipliers associated with moiety conservation constraints. The “stress” vector **t**_e_ = **y**_e_ − **h**_e_ contains the (total) fitness derivatives with respect to enzyme levels.

### 2.3 Metabolic balance equations

From the economic rules and assuming optimal enzyme levels, we obtain two balance equations for enzyme investments (see Figure 2). In enzyme-optimal states, the enzyme stress must vanish, **t**_e_ = 0, and by combing this with the economic rules (9), we obtain the balance equations (proof see SI section A)

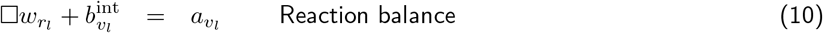

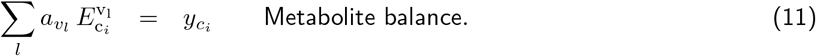

**Figure 2.**
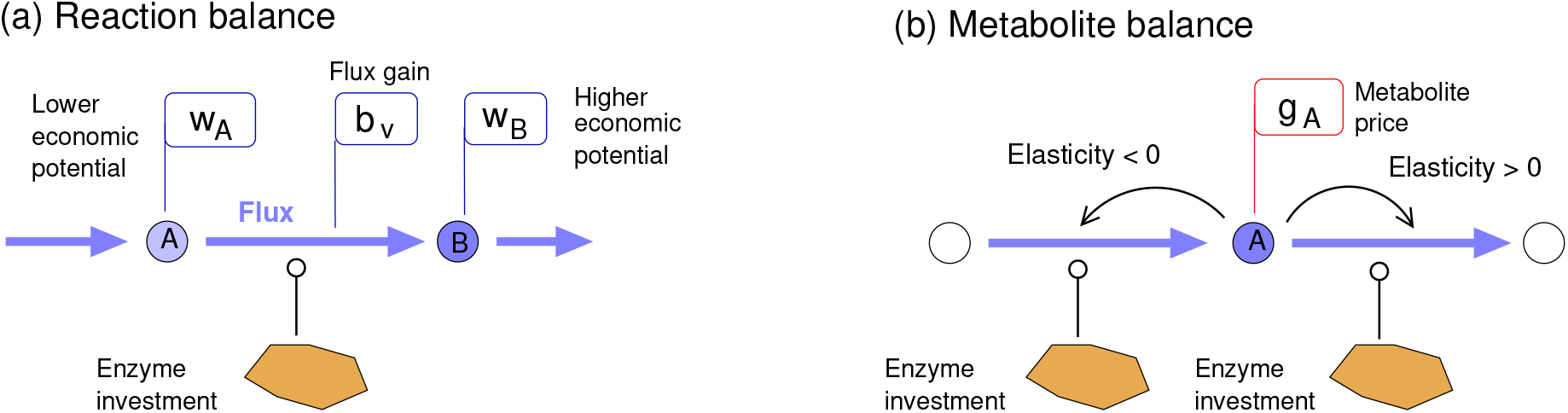
Economic balance equations. The equations for optimal states relate enzyme investments to economic value. (a) Reaction balance (10), **h**_e_*/***v** = □**w**_r_: in reactions without fitness direct effects, the enzyme investment must be balanced by value production, given by the economic potential difference multiplied by the flux. Polygons symbolize enzyme investments, similar to enzyme amounts in a proteomap (Fig. 1 (a)). (b) Metabolite balance (11), 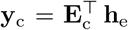. The balance equations hold for various modelling frameworks, including FBA-like models, kinetic models, and cell models.

Aside from production values (or “economic potentials”) 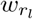, flux gains 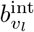, and economic loads 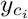, the equations contain the flux burdens 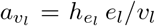. Equation (10) holds for kinetic and stoichiometric models in optimal states and follows directly from mass-balance constraints on metabolites, while Equation (11) results from enzyme kinetics. Like Eq. (4), the balance equations (10) and (11) are fully local: while the economic variables describe indirect (i.e. “network-wide”) fitness effects (of metabolite production and concentration), they are coupled only locally in the equations, i.e. between neighbouring network elements.

The economic values in Eq. (10) and (11) represent fitness derivatives (gains, prices, burden, and potentials). Since derivatives are taken with respect to different physical variables, the resulting economic variables show different units. If we multiply each of them by the dual physical variable (or equivalently, if we use logarithmic derivatives), we obtain new economic values called point cost (or “cost contribution”) and point benefit (or “benefit contribution”), which all have the same units, namely units of fitness. They satisfy the equations in “value production form”

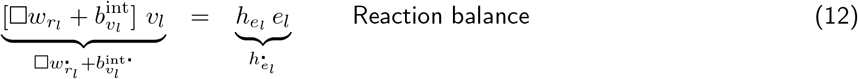

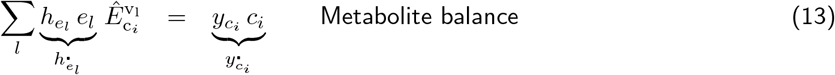

where 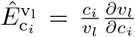 denotes scaled elasticity coefficients. The first equation is exactly our Equation (4). The balance equations relate physical model variables (fluxes *v*_*l*_, metabolite concentrations *c*_*i*_, and enzyme levels *e*_*l*_) to economic variables.

The balance equations (12) and (13) for metabolic systems in optimal states relate enzyme investments 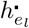 to network structure, kinetics, and fitness objectives. The reaction balance Eq. (12) equates an enzyme investment to the net value production in the catalysed reaction. Since enzyme investments are positive, every reaction must produce positive value. The flux value is given by the sum of economic potential difference and flux gain, and value production is given by the flux value multiplied by the flux. In reactions without direct flux gain, fluxes must run from lower to higher potentials to achieve a positive value production. This situation resembles reaction thermodynamics, where fluxes must run from higher to lower chemical potentials to dissipate Gibbs free energy. Without direct flux gains, the positive enzyme investments imply that the economic potentials increase in every reaction: in fact, every metabolite embodies in its economic potential, the cost of all the transporters or enzymes that were needed to import or produce this metabolite. Importantly, Eq. (12) applies not only to single reactions but also to entire pathways. A pathway flux can be defined by a flux vector **v**^pw^ that is stationary on the pathway and 0 outside the pathway. With this definition, the total pathway enzyme investment **h**_e_ · **e** must be equal to the value production rate in the pathway, 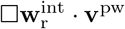.

Using the economic balance equations, known enzyme costs and benefits can be “propagated” across the network: if the total enzyme investment in a pathway is known, we also know what benefit the pathway must provide; if we further know the flux and the economic potentials of substrates “flowing in”, we can infer the economic potential of the product. Conversely, the fluxes and flux values in a pathway determine the enzyme investments required. We can compute them from the balance equations: Eq. (12) determines the sum of investments, and Eq. (13) determines their ratios. For example, in a linear metabolic pathway without metabolite bounds, the enzyme investments in producing and consuming reactions of a metabolite must be inversely proportional to the reaction elasticities. Known elasticities will tell us about the ratios of enzyme investments, and by tracing these ratios along the pathway, we obtain the relative investments in all pathway enzymes. In a model with all kinetic constants set to 1, the enzyme investments would decrease along the chain^4^, as has been shown for flux optimisation at a fixed total enzyme level [15].

## 3 Economic laws derived from metabolic optimality conditions

### 3.1 Economic variables defined through Lagrange multipliers

While I postulated that economic variables exist, I did not explain so far how they are defined and how their balance equations can be derived. My definition is based on existing metabolic optimality problems [13]: by writing such a problem in an “expanded form” with explicit constraints, we obtain the economic balance equations, where Lagrange multipliers serve as economic variables,.

Lagrange multipliers, a tool from constrained optimisation, play a key role in physical theories that are based on variational principles (such as Lagrangian mechanics and classical thermodynamics). Mathematically, they provide a way to handle constraints in optimality problems. Geometriclly, equality constraints define a manifold of feasible states, and in a local optimum the fitness gradient must be orthogonal on this manifold. This criterion can be formulated using Lagrange multipliers. With control variables in a vector **x**, a fitness function *f* (**x**), and equality constraints *g*_*i*_(**x**) = 0, in a local optimum state there will be numbers *α*_*i*_ (called Lagrange multipliers) such that ∇*f* + ∑_*i*_ *α*_*i*_ ∇*g*_*i*_ = 0 holds, where ∇ denotes the fitness gradients and *α*_*i*_ is the Lagrange multiplier related to the *i*^th^ constraint. For an optimal state to exist, there must be Lagrange multipliers *α*_*i*_ that solve this equation. The well-known optimality condition ∇*f* = 0 in problems without constraints is an important special case. We can also formulate this differently: based on the functions *f*(**x**) and **g**(**x**), we define the Lagrange function ℒ(**x, *α***) = *f* (**x**) + ***α***^*T*^**g**(**x**). Optimality (of *f*(**x**) under constraints **g**(**x**) = 0) requires that *∂*_**x**_ℒ(**x, *α***) = 0 for some choice of ***α***.

Lagrange multipliers can also handle inequality constraints. In problems with inequality constraints, optimal states must satisfy the Karush-Kuhn-Tucker (KKT) conditions (SI **??**). The Lagrange multipliers have particular signs, depending on the type of bound (lower or upper) and whether the bounds is active. In maximisation problems (the problems considered below), active lower bounds lead to positive signs and active upper bounds lead to negative signs (see SI section **??**), while inactive bounds lead to zeros (i.e. effectively, no shadow value at all). Optimality problems with inequality constraints appear, for example, in linear programming (LP) problems.

In summary, each constraint defines a Lagrange multiplier, and the optimality conditions are equations between Lagrange multipliers and derivatives of objective and constraint functions. The numerical value of a Lagrange multiplier in the optimal state is called shadow value or shadow price. Most linear programming solvers return shadow values as a standard output. In Flux Balance Analysis, shadow values arising from mass balance constraints have been used to detect valuable (e.g. growth-limiting) metabolites. On a more theoretical level, Warren and Jones showed that shadow values in FBA resemble chemical potentials in reaction thermodynamics [16].

Extending these works, below I define economic variables (or “metabolic values”) more generally by shadow values in given metabolic optimality problems. The aim is to associate for each physical variable (e.g. the “value of an enzyme concentration”) with an economic value, a new property of the corresponding model element (e.g. an enzyme). However, in defining economic values as shadow values, there is a conceptual problem: mathematically Lagrange multipliers are not associated with model variables, but with *constraints* (which may refer to one or several variables, and which may not exist for all variables). Therefore, I write the optimality problems in “expanded form”, a form in which all model variables are treated as control variables and all dependencies between them (e.g. rate laws yielding metabolic fluxes from compound concentrations) are written as explicit constraints. In this formulation, each constraint belongs to a physical variable and the Lagrange multipliers for the constraint can be seen as a “dual” of that physical variable.

In the rest of this section, I show how the economic laws can be derived from different metabolic optimality problems (see Figure 3 for an overview). In each case, I write down their optimality conditions and translate each of them into economic balance equations. The underlying metabolic models describe enzyme and metabolite concentrations, fluxes and production rates. Concentrations and fluxes are coupled by two types of laws: concentrations determine fluxes via rate laws, and fluxes determine concentration changes in time, which are set to zero to ensure a steady state. All these problems – in flux, metabolite, or enzyme space – are simplified versions of a general optimality problem on the maifold of possible metabolic states [13]. For convenience, I write them as *maximisation* problems. In this case, lower bounds yield positive shadow values (resembling benefit terms), while upper bounds yield negative shadow values (resembling cost terms) – (see SI **??**). For detailed descriptions, see SI section A. The derivations will be repetitive, because deriving the economic laws follows always the same procedure. Later we will see a general formulation that avoids repetitions and highlights similarities between all these approaches.

**Figure 3.**
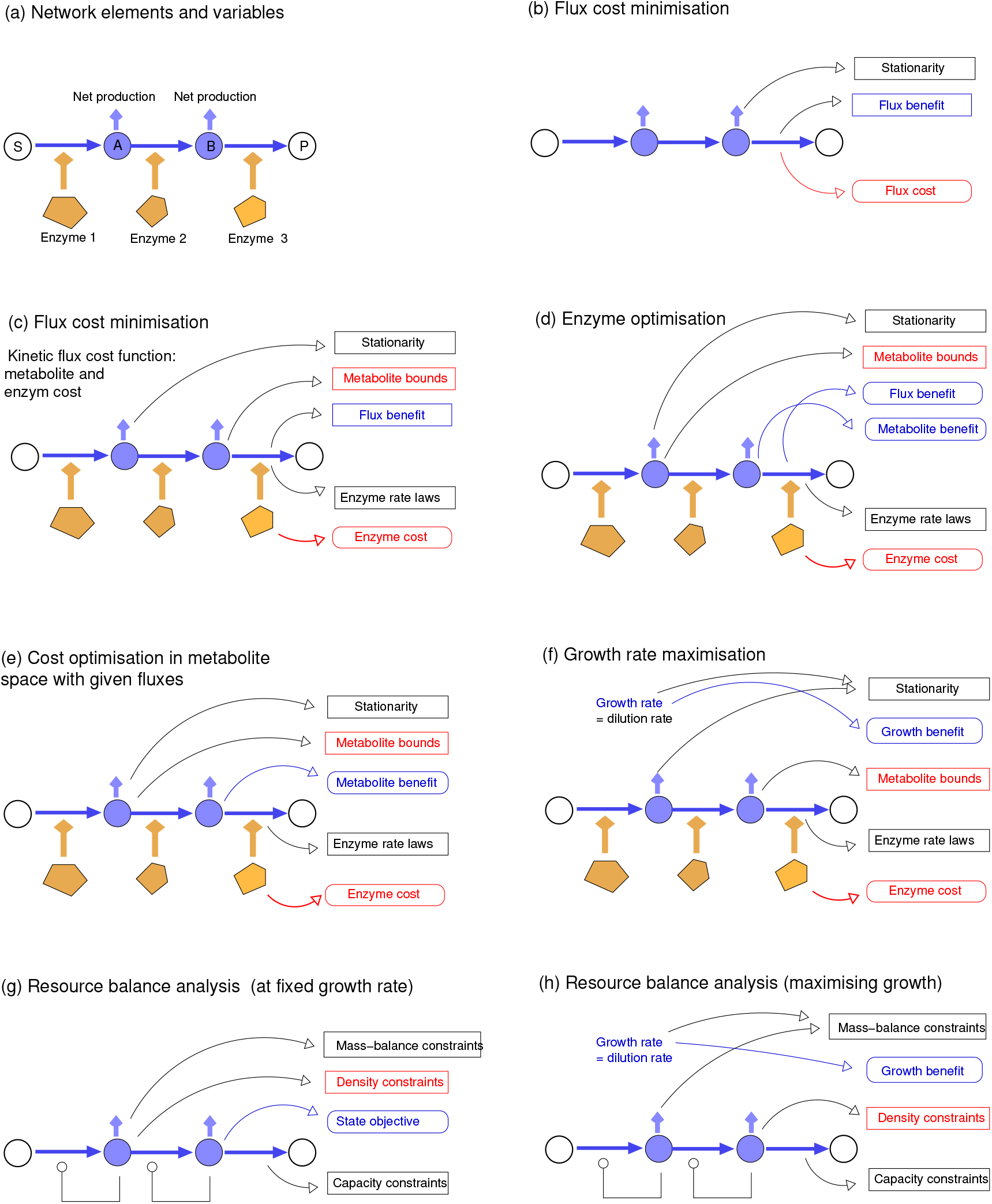
Metabolic optimality problems described in the paper. (a) Example pathway. The network elements (metabolites, reactions, and enzymes) carry physical variables (concentrations, fluxes or production rates) which are constrained (e.g. by stationarity or density constraints) and scored by target functions (cost and benefit terms). By optimising or constraining the target variables, different types of optimality problems can be obtained. (b) Flux cost minimisation. Boxes show constraints (boxes) and objectives (rounded boxes); benefit terms and lower bounds shown in blue, cost terms and upper bounds shown in red. For clarity, only some of the arrows are shown. For example, stationarity (net production rate = 0) holds for all internal metabolites, but is depicted only for metabolite B. (c) Flux cost minimisation. A “kinetic flux cost function” represents metabolite and enzyme cost. (d) Enzyme optimisation in a kinetic model. (e) Cost optimisation in metabolite space (“Enzyme cost minimisation”). (f) Maximising the cell growth rate. (g) Resource balance analysis problem with a linear objective, scoring fluxes and compound concentrations. Compounds (blue) comprise metabolites and proteins. To obtain a tractable problem, nonlinear rate laws are replaced by capacity constraints, linear in the catalyst level and independent of reactant concentrations. (h) Resource balance analysis problem for maximal growth rate.

**Figure 4.**
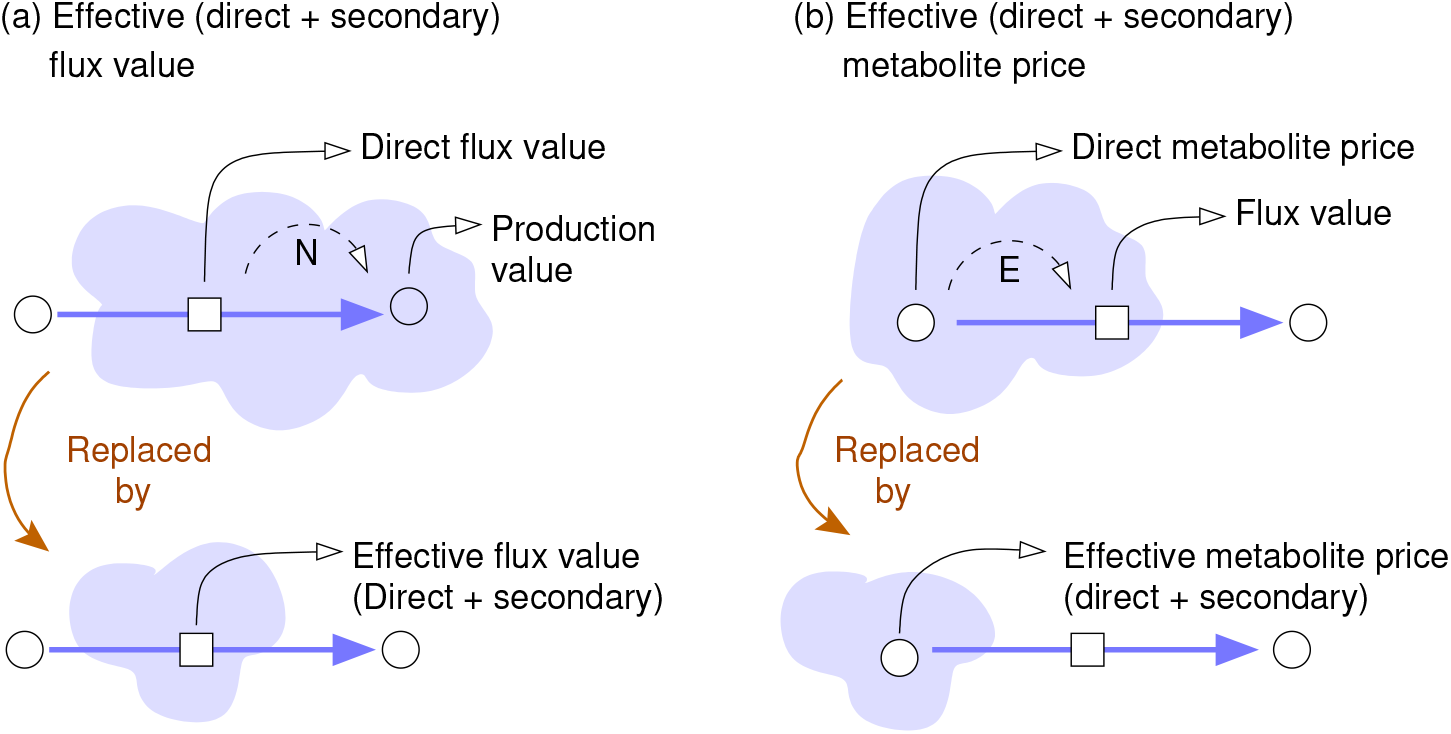
Direct, indirect, and total values.

### 3.2 Flux analysis models

Our first metabolic problem is Flux Cost Minimisation (FCM): we search for a flux profile **v** that realises a given linear^5^ benefit 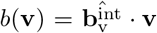 at a minimal cost *a*(**v**) (see Figure 3 (b) and (c), and SI section **??**). The cost function *a*(**v**) may be chosen ad hoc from models (e.g. describing the enzyme cost in an underlying kinetic model). Compound concentrations and kinetics are not considered. An example of this approach is FBA with minimal fluxes [17]. The FCM problem reads

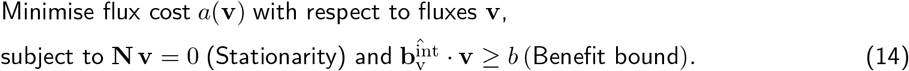

The stoichiometric matrix **N** refers to internal (i.e. balanced) metabolites. Typically, the benefit function *b* describes the production rate of a valuable product or biomass, while the cost function can be a weighted sum of the absolute fluxes or a general non-linear function. For a plausible model, fluxes should be costly, i.e. the cost *a*(**v**) must increase with the absolute flux (i.e. sign(*∂a/∂v*_*l*_) = sign(*v*_*l*_); “assumption of costly fluxes”). In the optimality problem (14), an inequality constraint *b*(**v**) *> b* ensures a desired benefit value. Since the constraint will always be active in optimal states, we could also use an equality constraint 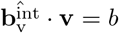, but we consider the inequality constraints for didactical reasons, to learn about the sign of the corresponding shadow value. To solve problem (14), we rewrite it as a maximisation problem, introduce Lagrange multipliers (a vector ***α*** for stationarity, and a scalar *β* for the flux benefit constraint), and define the Lagrangian ℒ. Now the optimality condition reads

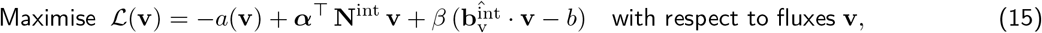

and with the constraints from (14). The optimality conditions (Karush-Kuhn-Tucker conditions, see SI section **??**) read

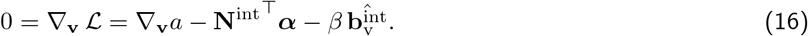

For a flux profile **v** to be optimal, there must be a vector ***α*** and a scalar *β* that solve this equation. Since (15) is a maximisation problem and *β* belongs to a lower bound, the KKT conditions require that *β* must be positive (see SI section **??**). Now, we introduce new variable names to write Equation (16) in a standard form comparable between optimality problems. By defining internal economic potentials 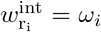, direct flux gains 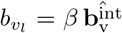, and flux prices 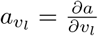, and noting that 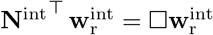, we obtain the reaction balance

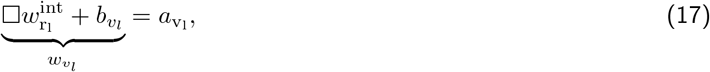

which must hold for every active reaction *l*. Remember that 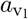 must have the same signs as *v*_*l*_. Importantly, Eq. (17) contains only local variables: the (local) flux price 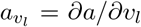 must be balanced by the (local) flux value 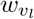, given by the sum of flux gain 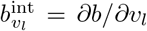 and economic potential difference 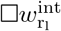 in the same reaction.. The balance equation holds for FCM problems and similar optimality problems. Interestingly, the same procedure can also be used to derive thermodynamic laws (see SI sections A and **??**). We assume that all metabolic fluxes must satisfy a principle of minimal entropy production and formulate this as an FCM problem: the resulting balance equations tell us that all fluxes must dissipate Gibbs free energy and must lead from higher to lower chemical potentials.

Fluxes are costly because of their enzyme demand. How can we write this cost as a flux cost? By multiplying Eq. (17) with the flux *v*_*l*_, we obtain the reaction balance in “value production form”

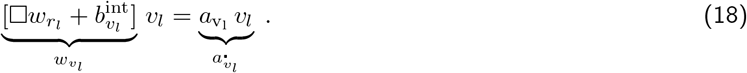

The *flux point cost* 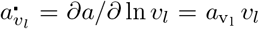, on the right, is derived from our flux cost function and measured in units of fitness (Darwins, Dw). We can now define a flux cost function that represents enzyme cost. Given the kinetic rate laws, which describe a relationship between enzyme levels, metabolite concentrations, and fluxes, the enzymatic cost *a*^enz^(**v**) of a flux profile **v** is defined as the minimal enzyme cost at which **v** can be realised with these rate laws and assuming optimal adjusted enzyme levels (see SI sections A and **??**). With an enzyme cost function *h*(**e**) and enzyme prices 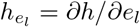 we obtain the reaction balance

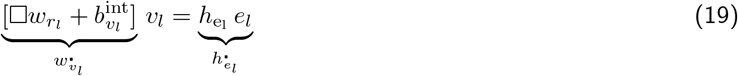

which relates flux values 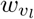 and fluxes **v** to the enzyme investment 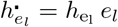. By comparing Eqs (18) and (19), we find that 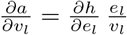 (proof in SI **??**). This means: flux investment (in an FCM problem with an enzymatic flux cost function *a*(**v**)) and enzyme investment (in the underlying kinetic model) are equal: 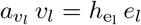.

### 3.3 Kinetic models

We saw that the mass balance constraints in FCM define shadow values that satisfy the reaction balance Equation (12) and can be seen as economic potentials. To obtain our second balance Equation (13), we now consider kinetic models with metabolite concentrations, enzyme levels, and fluxes as model variables (see Figure 3 (d), and SI section **??**). The variables must be chosen to maximise a metabolic fitness Eq. (1), comprising flux benefit, metabolite cost, and enzyme cost. With enzyme levels as choice variables^6^, defining a stable steady state^7^, we obtain the optimality problem

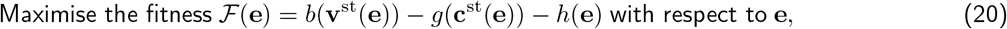

where steady-state fluxes **v**^st^(**e**) and metabolite concentrations **c**^st^(**e**), are implicitly defined by stationarity, rate laws, and moiety conservation:

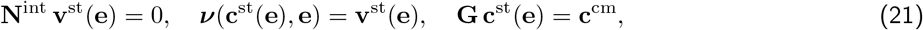

With the internal stoichiometric matrix **N**^int^, rate laws *v*_*l*_(*·*), conserved moiety conservation matrix **G**, and conserved moiety concentrations 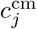. Since the functions **c**^st^(**e**) and **v**^st^(**e**) are hard to compute, we go back to Eq. (20), treating **e, c**, and **v** as free variables and use Eq. (21) as explicit constraints. In the resulting “expanded form”, the problem now reads

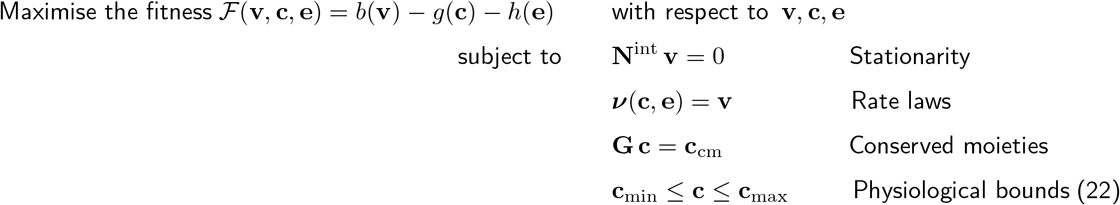

If we write down the optimality conditions, we obtain the economic rules in Eq. (9) (see SI sections A and **??**). By combining these rules, we further obtain two balance equations: our reaction balance for fluxes and metabolite production, and a metabolite balance for optimal metabolite concentations.

In a model with given fluxes, enzyme and metabolite concentrations can be varied and optimised for a minimal cost (see Figure 3 (e)). Maximising the fitness (1) at fixed fluxes yields to an optimality problem for metabolite and enzyme profiles. Given the fluxes in in **v** and the rate laws *v*_*l*_(**c, e**) = *e*_*l*_ *k*_*l*_(**c**), each metabolite profile **c** defines an enzyme demand 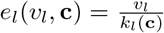. With this demand function, enzyme cost *h*(**e**) and metabolite cost *g*(**c**) can be combined into an effective “kinetic metabolite cost” *g*^kin^(**c**) = *g*(**c**) + *h*(**e**(**v, c**)) for metabolites, including an “enzyme overhead cost”. For given fluxes **v** and concentrations **c**, this cost is easy to calculate. We now fix a flux distribution **v** and treat metabolite concentrations as free variables: if we constrain them by physiological bounds^8^ and moiety concentrations, we obtain the optimality problem

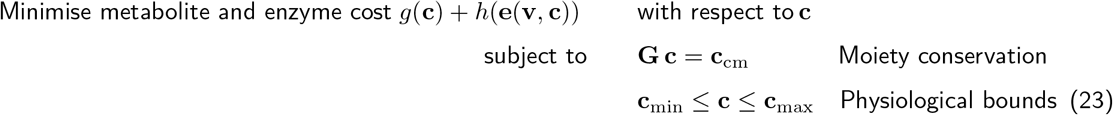

In expanded form (with metabolite concentrations **c** and enzyme levels **e** as control variables and dependencies as explicit constraints), our problem reads

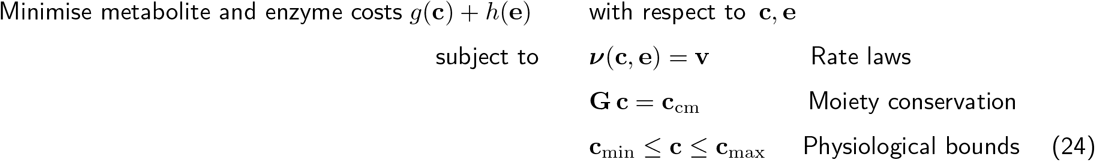

It resembles Eq. (22), but now with predefined fluxes (which makes flux benefits and stationarity constraints obsolete). From the optimality conditions with Lagrange multipliers (see SI sections A and **??**), we obtain once more the metabolite balance (11). This time, there are no economic potentials and no reaction balance equation: since the fluxes are predefined, there is no need to enforce stationarity by mass-balance constraints, and without the constraint there are no corresponding economic variables.

### 3.4 Resource allocation models

Living cells must be able to replicate all their components to ensure growth and self-repair. Aside from membranes and DNA, which define the cell, there need to be molecular machines to produce these components, other machinery to produce these machines, and so on. In short, for each cell component there must also be the machinery to produce (or to import) it, a self-consistency requirement for living beings dubbed the “closure to efficient causation” by Robert Rosen [18, 19]. Closure to efficient causation does not only concern the inventory of a cell, but also amounts! At balanced growth, all compound concentrations and production rates must be fine-tuned: each compound must be reproduced at the right rate to balance its dilution. For each cell component with concentration *c*_*i*_, we need a net production rate *r*_*i*_ = *λ c*_*i*_ and the right concentration of the producing machines (which depends on their catalytic rates). This requires a sufficient production of other machines, and so on. Finally, since all compounds occupy space, their concentrations (or abundances) are limited by density constraints. Until here, the cell state is not uniquely defined; concentrations and fluxes are coupled through various dependencies and constraints, but there may still be various possible arrangements for a viable growing cell. At high growth rates, these conditions become harder to satisfy, and above some critical growth rate, it is impossible to satisfy all the constraints. This eventually limits the possible growth rates. To maximise growth computationally, we need to find the maximal possible *λ*. A growth maximisation under constraints has been considered in various modelling frameworks, including small or large, and kinetic or stoichiometric models of cells.

A main difference between metabolic models and cell models is how they treat cell growth and the dilution of compounds. In models of self-replicating cells, growth itself may be the maximisation objective, which means that fast dilution is “good”. In metabolic models, dilution is usually neglected and a “biomass production” objective is used, i.e., the synthesis rate of precursors for macromolecule production (which itself is not modelled).

Resource Balance Analysis (RBA) implements these ideas to build detailed resource allocation models of cells (SI section **??**). While RBA models may be large, comprising thousands of variables, the basic formulae are rather simple. We describe all compounds (metabolites, macromolecules, and molecular complexes) by a concentration vector **c** and all reactions and macromolecular processes by a flux vector **v** (see Figure 3 (g) and (h)). The two vectors are related by a stoichiometric matrix **N**^int^. To obtain a tractable problem, all nonlinearities are removed. Instead of the nonlinear rate laws (depending on **e** and **c**), we assume for each enzyme (or macromolecular machine) a fixed catalytic efficiency (i.e. catalysed flux per catalyst concentration) that is independent of metabolite concentrations and given by a model parameter. For reversible reactions, we assume that reaction rates can assume any value between their maximal velocities 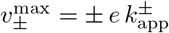 in forward and reverse direction (which in turn depend linearly on the enzyme level, and on nothing else). The catalytic rates for all reactions are given in matrices **E**_for_ (forward enzyme elasticities) and **E**_rev_ (reverse enzyme elasticities). Density constraints limit a weighted sum of protein levels, and the weights are given by a (non-negative) matrix **D**. We obtain three types of constraints:

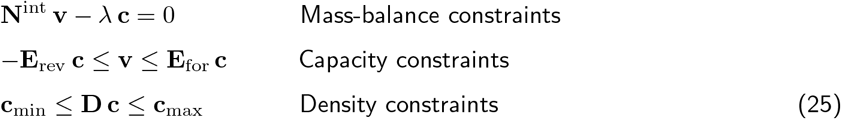

Given a growth rate *λ*, the constraints define a feasible region for **c** and **v**. Above some growth rate *λ*^max^, this region has a zero volume and the constraints cannot be satisfied. For lower growth rates *λ*, the constraints defaine a feasible polytope for the state vector (**c, v**). To describe whether a given *λ* allows for a feasible state, a linear satisfiability problem must be solved. Within the set of feasible states defined by these constraints, we can consider different optimality problems.

In the first RBA problem, a cell must grow at a given (cell-specific) rate *λ*, while its state variables **v** and **c** must maximise some linear objective 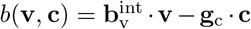 under the constraints Eq. (25). This is a linear problem, and LP solvers return the shadow values directly. There are shadow values 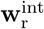 associated with mass-balance constraints, **a**_v_, associated with capacity constraints, and **g**_d_ associated with density constraints and with an effective compound price vector 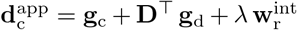. These shadow values satisfy the economic balance equations (see appendix A)

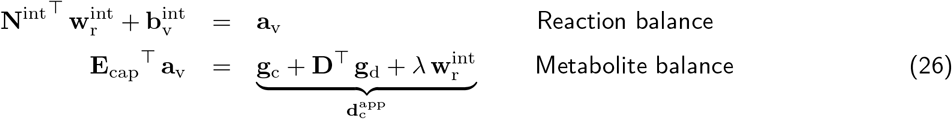

The matrix **E**_cap_ contains the actual catalytic rates, i.e. the catalytic rates in forward or reverse direction, corresponding to the flux direction in the optimal state. As before, flux burdens 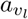 and fluxes *v*_*l*_ must have the same signs (to ensure positive enzyme investments), and **g**_d_ must be positive (except for inactive density constraints, which lead to vanishing components 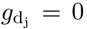). Thus, the shadow values in our RBA problem can be seen as economic potentials; with a linear objective, these are the shadow values returned by the LP solver!

In our second RBA problem, growth itself is the fitness objective. To determine the maximal growth rate *λ*^max^ and the corresponding cell state, we maximise *λ* under the constraints (25) and treat *λ*, **v**, and **c** as control variables. This would be a nonlinear problem, but since we know that above some *λ*^max^ the problem is infeasible, we treat *λ* as a parameter and decide, for each *λ*, whether (25) is solvable, and approximate *λ*^max^ by a dichotomy search. This time, the solver will not return shadow values for our objective (the growth rate). However, we may still obtain them from the economic balance equations! The optimality conditions for maximal growth under the constraints (25) lead to economic balance equations for internal economic potentials 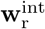, flux burdens **a**_v_, and “density prices” **g**_d_ (derivation see Eq. (34) in appendix)

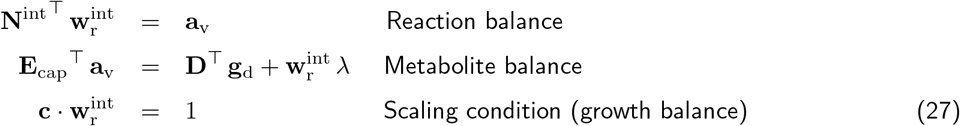

Instead of maximising *λ*, we may also optimise other model parameters at a fixed growth rate *λ*. For example, for growth on glucose we may search for the minimal external glucose concentration at which a cell can survive: we can do this by minimising the glucose transporter efficiency – the catalytic rate *E*_for_ of this transporter, which in reality depends on the external glucose level – as a model parameter. Unlike optimising a model variable (or a linear function of them), this is not an LP problem. We may again resort to a dichotomy search, but we first need to show, like for the growth rate, that there is a threshold parameter value above which (and only above which) the model is feasible. Once this has been proven, we can proceed like in growth rate maximisation.

### 3.5 Metabolic oscillations

Metabolic value theory can be applied to temporal dynamics. As a simple case, we consider driven oscillations in metabolic concentrations and fluxes caused by periodic enzyme levels (and possibly, external metabolite con-centrations). To model such oscillations, we linearise the model dynamics and describe enzyme levels, metabolite concentrations, and fluxes by sine-wave profiles. The complex-valued amplitudes (with amplitude vectors 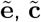 and 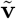) depend on the circular frequency *ω* = 2 *π f*, and their dynamics is determined by mass balances 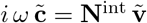 and linearised rate laws 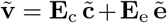. In optimality problems for enzyme oscillations, periodic states are scored by a fitness function, and the enzyme amplitudes and phase angles are optimised for a maximal fitness (see SI section **??** and [20, 21] for formulae).

Mathematically, metabolic oscillations resemble metabolic states with dilution. The mass-balance equation 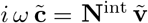 resembles the stationarity condition 0 = **N**^int^ **v** − *λ* **c** for steady states with dilution. Thus, to model metabolic oscillations, we may consider a model with dilution, write down the steady-state condition and replace the dilution rate by an imaginary frequency *λ* → *iω* (with real-valued circular frequency *ω*): this yields the mass-balance equation for oscillatory states^9^! The analogy between dilution and oscillations can be useful: in modelling, methods for models with dilution can be reused to describe oscillations.

## 4 Metabolic value theory

### 4.1 Why do economic balance equations hold generally?

In our previous examples, all economic laws were derived in the same way: we convert a model into expanded form, write down the optimality conditions with Lagrange multipliers, and arrange the terms into “direct” and “indirect” effects. These terms are then called economic variables. The same recipe works for a wide range of models (with various types of variables, constraints, and objectives). The steps from an optimality problem to economic balance equations are shown in Box 1. But why is it that all these optimality problems yield the same economic balance equations? If some models consider kinetics and other are purely stoichiometric, how can we obtain exactly the same balance equations? The reason is that the same type of constraints will always lead to the same economic laws. Let us consider different models of the same system, describing the same physical reality: they rely on the same metabolic network, the same rate laws (if kinetics are considered), and the same metabolic targets. The fact that the models assume the same constraints between physical variables leads to economic laws of the same form. For example, FBA and kinetic steady-state models apply the same mass-balance constraints. If we write them in expanded form, then in both cases we obtain economic potentials as the associated shadow values. No matter whether we optimise or constrain a metabolic target variable, we obtain the same types of terms in the balance equations. This means: the model formulation may change, but the shape of the economic balance equation remains! This allows us to compare economic variables between models, even if models contain different types of variables (only fluxes in FBA, fluxes and concentrations in kinetic models) and treat costs and benefits differently (as objectives or constraints). Since our economic laws hold for various model types, we can claim that they are not properties of specific models, but describe optimal metabolic states themselves, thus capturing facts in reality. If economic variables and economic laws are model-independent, there is certainly space for a general theory.

### 4.2 Metabolic optimality problems in expanded form

A metabolic model^10^ describes compounds (with concentrations) and reactions (with fluxes). The model variables are typically bounded (constrained to physiological ranges) and connected by three types of constraints: (i) mass balance constraints, relating fluxes to metabolite net rates, and assuming that internal net rates vanish; (ii) kinetic rate laws or capacity constraints, relating enzymes concentrations to reaction fluxes; and (iii) density constraints delimiting total concentrations in cell compartments and membranes.

Below, we often consider kinetic models in a “standard form” with variables **c, e, v**, and **r** (see Box 2). The variables are interrelated (metabolite rates *r*_*i*_ depend on reaction rate *v*_*l*_, and reaction rates *v*_*l*_ depend on con-centrations *c*_*i*_ and *e*_*l*_) and are restricted by fixed values or bounds (e.g. for internal metabolite rates). To define a metabolic optimality problem, we can start from a metabolic model and introduce cost and benefit functions (or “target functions”) to describe fitness and functional constraints. These functions are added as nodes to our dependence network. By convention, in our standard models fluxes are scored by a benefit function, while metabolite and enzyme profiles are scored by cost functions. In our “standard” models, we simply maximise a fitness function given by the benefit-cost difference *𝓕*(**v, c, e**) = *b*(**v**) − *g*(**c**) − *h*(**e**), but there are many other possibilities: we may consider other combinations (e.g. defining fitness as a function of the production/enzyme ratio *b*(**v**)*/h*(**e**)), a constrained optimisation (e.g. maximizing production *b*(**v**) at a fixed enzyme budget *h*(**e**)), or a Pareto optimisation (between two or several targets). In all these cases, the optimality conditions concern linear combinations of the target functions’ gradients in the space of our model variables.

#### Box 2

Economic balance equations derived step by step

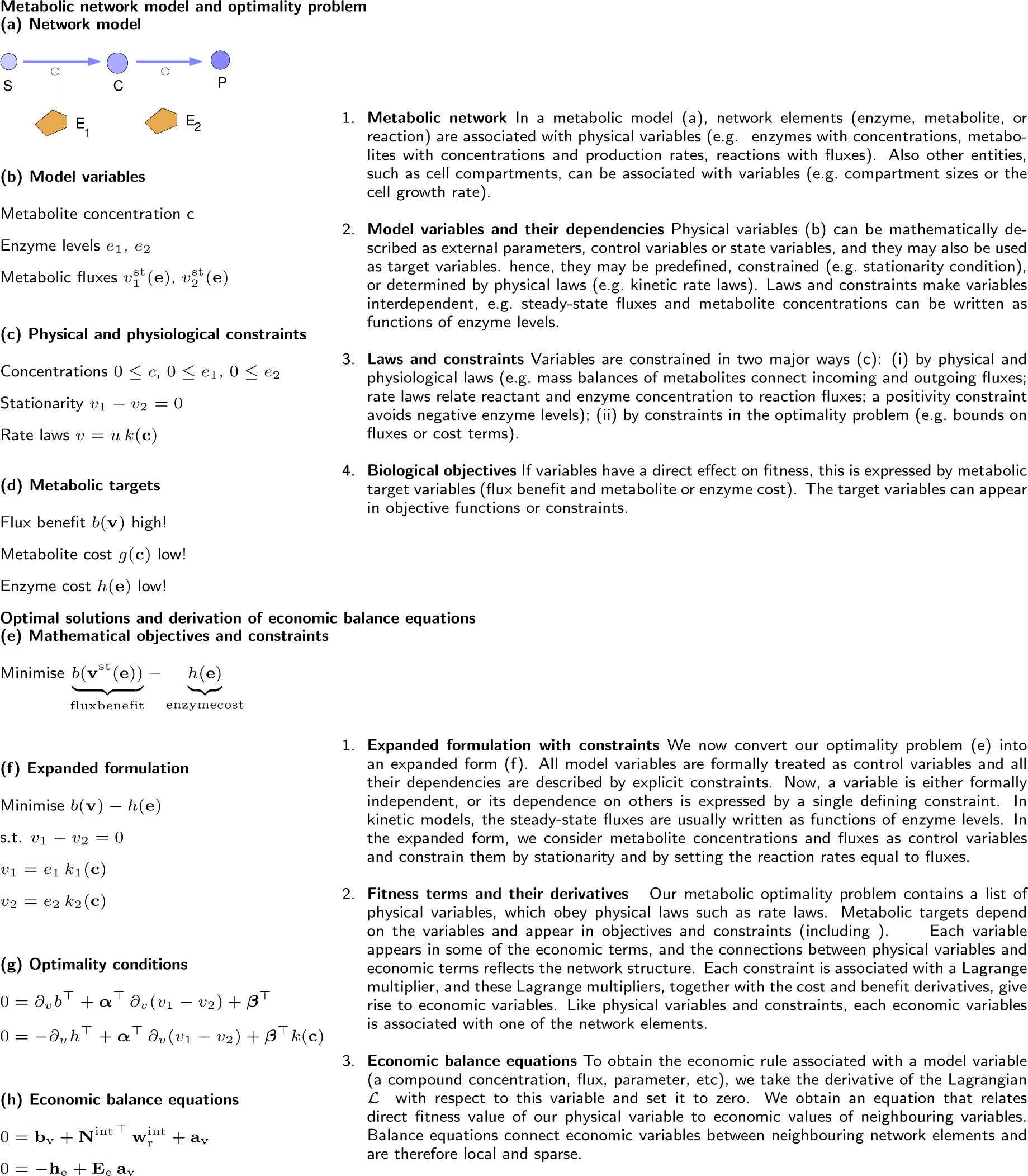

#### Box 3

Economic variables and economic rules

In Metabolic Value Theory, each physical model variable *x* is associated with an economic value *f*_*x*_ (e.g. each a reaction flux is associated with a flux value). The total economic value *f*_**x**_ describes the effective fitness effect *δf* = *f*_**x**_ *δx* of a variation *δx*. This fitness effect can consist of several contributions. First, a variable may have a direct effect on the fitness function, described by a partial derivative *∂f/∂***x**. Second, *δ***x** may violate lower or upper bounds of individual variables, associated with shadow values. These two effects can be merged into a “direct fitness effect” (called gain or price). Third, a variable has effects on neighbouring child variables (see Box 2). If these child variables have (direct or indirect) effects on fitness, our variable “acquires” from them an indirect value. Formally, economic values can be defined by associating each physical variable with a constraint (defining this variable as a function of others, or equating it to a given external or control variable). The constraint gives rise to a shadow value, our economic value, which is dual to the physical variable and belongs to the same network element.

The following table lists the connection rules that relate the indirect value of a network element to the total values of “child” (i.e. elements whose associated variables are directly causally dependent on our focal variable) elements.

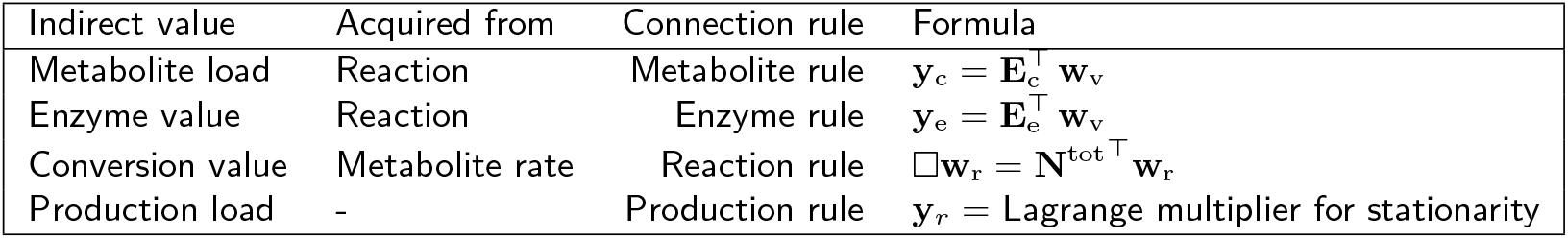

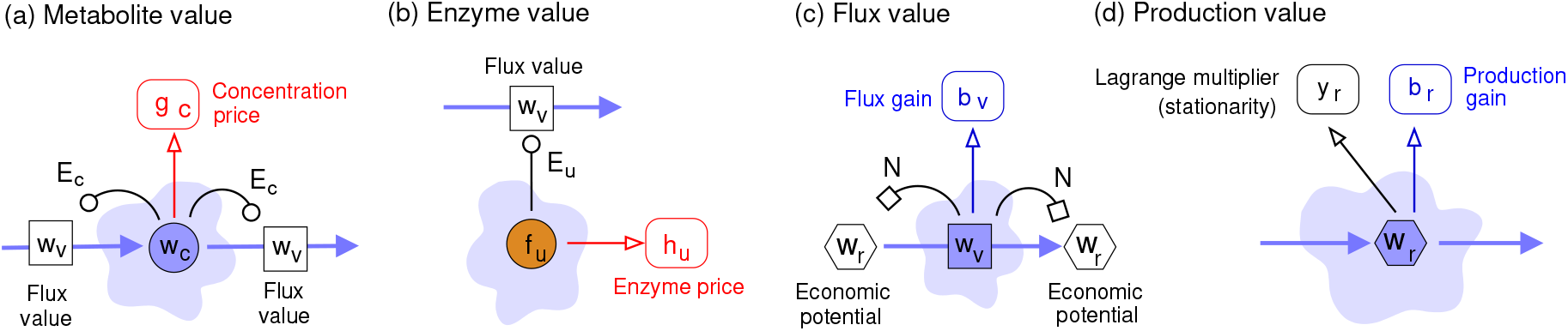

The economic rules state that economic values consist of a direct value (related to direct fitness contributions or active bounds) and an indirect value acquired from child elements (as described by connection rules.

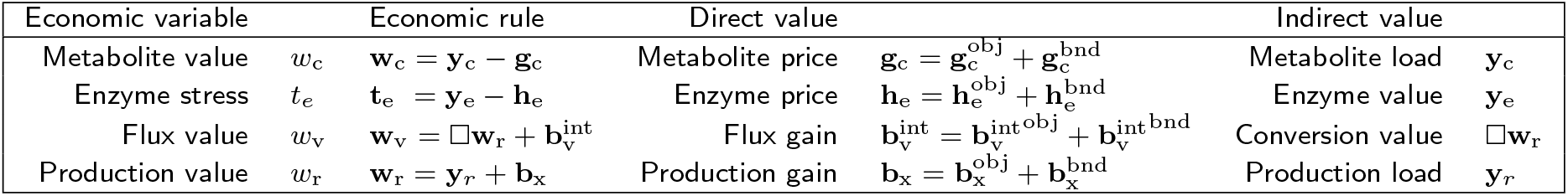

In optimality problem, the dependencies between model variables can be expressed in different ways: either, some variables are treated as control variables and the others are functions on them, or *all* variables are treated as control variables and all dependencies are written as explicit constraints. This second, “expanded” form is the form we choose here to formulate economic value theory more generally: all quantities appear as control variables, all dependencies are written as explicit constraints, determining a single variable each, and these dependencies must be acyclic (i.e. variables must be ordered and must only depend on “earlier” variables). This means that all physical variables are clearly separated, all constraints are known, and the benefit, cost, and constraint terms can be listed explicitly. Typical constraints in such models comprise (i) stationarity, i.e. the mass balance of internal compounds; (ii) given production and consumption rates of external compounds; (iii) rate equations, capacity constraints, and other physical laws (e.g. including density constraints or relations between compound concentrations and osmotic pressure and cell volume changes); (iv) bounds or predefined values for model parameters.

If we look at this more generally, we can consider optimality problems on general networks. In such problems, each model variable is associated with a network element, and we require that all constraints on variables are local, that is, they concern either individual single variables or neighbouring variables.

The derivatives of our target functions are called gains or prices. A non-zero gain or price of a physical variable *x* shows that *x* has a direct fitness effect. In our metabolic targets, we do not assume concentration prices for external metabolites (because their concentrations are predefined and cannot be altered), or production gains for internal metabolites (because their production rates vanish)^11^.

### 4.3 Economic variables and economic laws

A model variable can have direct and indirect fitness effects: if a variable *x* changes, this may affect fitness directly (if *x* appears directly as an argument in the fitness function), or it may affect other variables indirectly, which then have fitness effects. In any case, if we turn the causal relation (variable → fitness) around, we obtain an “incentive relation” (fitness → variable), which assigns a “use value” to our physical variable. For a simple example, let us see a chain of direct causal effects *x* → *y* → *z* → *f* : here fitness can be written as a function *f* (*z*(*y*(*x*))) and causal dependencies (for small variations) follow a chain rule: 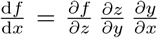. If we read this from right to left, we see how changes in *x* will change *f*. But if we read it from left to right, we see how *f* gives a “use value” to *z*, then to *y*, and so on; so use value is propagated in the direction *f* → *z* → *y* → *x*, opposite to the causal effects. The first term, 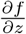, describes the direct value of *z*. The following terms 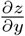 and 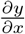 are “connections” by which a variable acquires a value from its child variable, and the entire formula yields the indirect value of **x** which emerges from a propagation of value (from *z* to *x*) backwards along the chain.

This was only a simple example. In general, our metabolic models are more complicated and we need to consider not direct causal effects, but steady-state changes. In a general description, each physical model variable (flux, metabolite net rate, metabolite concentration, enzyme level, and others) is associated with an economic dual variable or “economic value”, which describes a mapping from variations of the physical variable to variations in fitness. For infinitesimal variations (described by differentials *δ***x**), this mapping is linear, *δf* = **f**_**x**_ *· δ***x**, and the dual vector ^12^. **f**_**x**_ contains the direct economic values. However, there is a twist to this: since our model variables **x** are dependent, the variations *δ***x** are constrained to be “valid”, that is, respect all model constraints. With these constraints, we obtain Lagrange multipliers and effective dual variables that describe the indirect economic values (see Appendix B).

If a physical variable *u* is a control variable (e.g. an enzyme level) or an external variable (e.g. an external metabolite concentration) and **z**(*u*) is the metabolic steady state, the corresponding economic value is defined by the total derivative d*f/*d*u* of our fitness function with respect to *u*. Here is an example. The vector **y**_e_ = ∇_**e**_*f* (**z**(**e**)) maps variations *δ***e** of enzyme levels onto the resulting benefit changes *δq* = **y**_e_ *· δ***e**. The dual variables (or “economic values”) 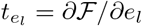, called enzyme stresses, form the fitness gradient in enzyme space and appear in the condition for enzyme-optimal states 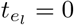. In contrast, if a physical variable *x* is one of the state variables (e.g. an internal production rate, flux, or internal metabolite concentration), we cannot vary it in the same way. To define the fitness derivative in this case, we introduce a perturbation parameter and compute fitness derivatives with respect to this parameter. Moreover, if a physical variable hits a constraint, an extra shadow value will be included in the economic variable.

In our network models in expanded form, each economic variable is related to a physical variable (and to its associated network element). For example, an internal metabolite (network element) has a net production rate (physical variable); accordingly, the corresponding economic value is called “production value” of this metabolite (or, for convenience, “economic potential”).

In the metabolic model, model variables are connected by laws (in the form of constraints). If a variables changes, this will change other variables (causally downstream), and if a variable is constrained, this may constrain other variables (causally upstream). Due to these connections, also variables that are not directly fitness-relevant may have an indirect impact on fitness.just liek physical variations are propagated between neighbour variables, also the values (i.e. the fitness incentives for variations) are propagated between neighbour variables (but in the opposite direction).

Above we saw how such variables can be defined: we derived it from optimality conditions with Lagrange multipliers. Alternatively, we may consider thought experiments with compensated variations, in which a flux is slightly increased, but its effects on the rest of the network are compensated by extra in- and outfluxes of its substrates and products. Finally, the same economic variables and laws can also be obtain from metabolic control coefficients.

### 4.4 Economic laws

In models in expanded form, direct dependencies between physical variables give rise to connection matrices (containing the direct sensitivities, and describing the direct propagation of infinitesimal perturbations). The same connection matrices also appear in economic rules. Fluxes and metabolite net rates, for example, are connected through the stoichiometric matrix (one type of connection matrix): *δ***r** = **N**^int^ *δ***v**. This is reflected in the economic variables: (indirect) flux values and economic potentials, too, are connected by 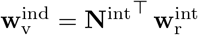. This holds more generally: physical variables and their economic counterparts are interlinked by the same connection matrices. This is how network structure (in the form of stoichiometric coefficients) and kinetics (in the form of elasticities) enter the economic rules Eq. (9).

It is a central insight from metabolic value theory that flux values can be written as local differences of production values (that is, “economic potentials”). Whatever indirect costs and benefit a reaction flux causes by its effects on the steady state, and wherever these costs and benefits may appear in the network, they can be described by saying that the reaction flux, locally, turns “cheaper” substrate into “more valuable” product: by assigning economic values to the inflowing substrate and outflowing product, we can fully capture (and replace in our formulae) the long-term, long-range effects of our reaction. This “replacement”, a central pillar of metabolic control theory, may sound astonishing at first. It actually means that the network-wide optimality problem, including the costs and benefits of all fluxes, metabolites, and enzymes, as well as all details of enzyme kinetics, can be encapsulated in simple local variables, just like the price of bread in a store encapsulates the complexity of our world-wide economic system.

Our examples and our general economic laws Eq. (44)? show that a model variable can contribute to metabolic economics in four different ways, which we describe by four different types of “effects”: (i) direct fitness effects, if a variable appears in the fitness function (in the diagrams, these effects are shown by single solid arrows); (ii) indirect fitness effects, if a variable causes steady-state changes with direct fitness effects elsewhere (this includes, for example, fitness effects due to moiety conservation; (iii) direct shadow values, caused by constraints on a single variable (e.g. bounds or positivity constraints); (iv) “stress” terms in non-optimal states, showing that our variable is not chosen optimally. In non-optimal states the total value of control variables does not vanish, but a “stress” remains. If we ignore this stress, the economic balance equations for non-optimal state become inequalities).

An optimality problem can be solved numerically to obtain the physical variables in the optimal state. But what about the economic variables? In LP problems, besides the physical most solvers also return shadow values, i.e. our economic values. In other problems, we can compute economic values numerically by varying model parameters (or perturbation parameters for system variables). in some cases, we may also compute them by solving the balance equations. This requires that all other terms in the equations be known, including connection matrices and direct values.

Economic variables and laws can be used to construct kinetic models in optimal states. can be systematically constructed. We start from network structure and an economical flux profile (we already know that uneconomical flux profiles cannot appear in optimal states). For the next step, a possible idea would be to choose reaction elasticities (e.g. by sampling [22]) and solve for the economic potentials. However, elasticities and economic potentials chosen at will may not yield an optimal state, and our formulae might lead to negative enzyme investments. To avoid this, we proceed differently: we first choose feasible economical fluxes and economic potentials satisfying the reaction balance equation. Then we construct economic loads and elasticities that satisfy the metabolite balance, and from the reaction elasticities we compute the kinetic constants. The resulting kinetic model will realise an enzyme-balanced state. However, one problem remains. In our algorithm, when choosing our economic potentials *ad hoc* (in step 1), we do not account for the predefined concentration price vector **g**_c_. Therefore, the economic potentials may not be compatible with this vector, and it may later turn out that **g**_c_ cannot be realised (in step 2). If this happens, we search for an approximate solution with a vector **g**_c_ *close to* the predefined one (e.g. requiring a feasible solution at a minimal adjustment of **g**_c_). Hence, in the construction **b**_v_ can be predefined, but **g**_c_ can be predefined only approximately.

### 4.5 Cost and benefit contributions and value flows

From our laws for economic values, we obtain similar laws for state variations. A state variation describes an (infinitesimal or finite) change of some of the model variables. In an optimal metabolic state, any legal (i.e. constraint-respecting), infinitesimal state variation must be fitness neutral. Otherwise, the variation (possibly with a minus sign) coudl be added to the state to further imporve the state, contrary to our assumption of an initial optimal state. Each law can be written in different forms, for values, point benefits, variations, or finite changes. Let us see an example. The economic rule 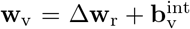 describes how the economic value of a flux arises from a difference of economic potentials and from a direct flux gain. This rule tells us about the fitness effects of variations. A (possibly non-steady) flux variation *δ***v** causes a variation of the metabolite net rates *δ***r** = **N**^int^ *δ***v**. By multiplying our rule by *δ***v**, we obtain a balance of differential fitness effects 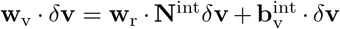, which can also be written as 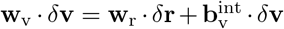. By integrating over **v**, we obtain the same formula, but for finite variations 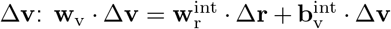. Finally, by choosing a Δ**v** = **v**, we obtain the balance 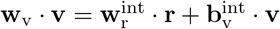 or in short 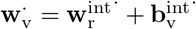. Altogether, we converted our economic rule from the original “value form” into a “value production form”, in which each economic value is multiplied with its physical dual variable. The same economic law, expressed in different forms, can tell us about values (fitness derivatives), differential and finite variations, and point benefits (fitness log-derivatives). The same procedure works for all economic rules and laws. Let us see this for another equation, our reaction balance Eq. (10). By multiplying with **v, v***/***e**, or *δ***v**, we obtain four alternative formulations:

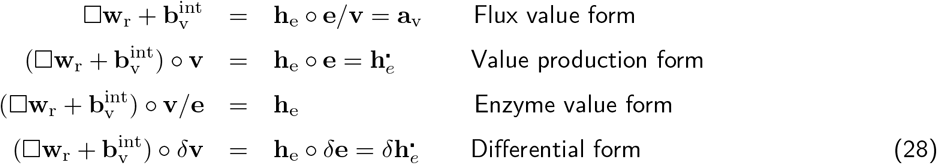

All four equations are equivalent.

The economic rules and balance equations in their “value production form” have an interesting interpretation. All terms – cost contributions (investments) and benefit contributions (value production) – have the same units, suggesting that they may be interconvertible. In fact, each of the rules can also be read as a conservation law, resembling continuity equations in physics that describe the flow of substances, charges, or energy. Like a physical substance, value enters the network in different places in the form of substrate or enzyme investments, it accumulates along the metabolic fluxes, and leaves the system where benefit is created (in the form of flux point benefits) or possibly, where fluxes or concentrations hit a bound^13^. If we say that “value is conserved”, this means that value can be transformed and can “move” between network elements, but the total amount of value cannot change: in an optimal state, whenever value flows into a network region (in the form of investments), it must leave the region again in the form of benefit contributions. In every part of the network, value inflows and outflows are balanced. Importantly, value conservation holds only in optimal states! In non-optimal states, “point stresss” (quantifying non-optimality) would appear in the balance equation, indicating a non-conservation of value^14^.

## 5 Discussion

### 5.1 A theory of metabolic value

How should a cell allocate its resources – such as proteins, energy, or cell space – to cellular functions, and how should it adapt fluxes and protein levels to changing conditions and tasks? Managing opposing needs at limited resources is a central question of economics. An economic perspective – asking not only what cells do in fact, but what they could do and how they are doing – can cast new light on metabolic behaviour. In economic models of metabolism, we typically formulate our questions as satisfiability or optimality problems, assuming that cell can “choose” between different configurations allowed by physics, and may do so by optimising (or finding best compromises between) simple fitness objectives. Biological optimality principles continue a tradition of variational principles in physics, engineering, and economics. In physics, variational principles^15^ provide a convenient formulation of many physical laws, describing for example laws for the trajectories of particles or light rays, or the configurations of electromagnetic fields.

We saw that optimality problems give rise to a “value structure”, a pattern of values and prices or benefits and costs on the metabolic network, which complements metabolic dynamics (and may partially explain it). This value structure resembles the pattern of chemical potentials and driving forces in thermodynamics, and like the thermodynamic variables, it shapes metabolic behaviour. We also saw how the value structure – the pattern of economic variables in the network – can be determined. In [13], I had described optimality problems for metabolic states (**v, c, e**) can be formulaed on a state manifold and be reformulated as problems for fluxes, metabolite concentrations, or enzyme levels alone. The predicted optimal states have some remarkable properties: futile flux patterns are suppressed and active enzyme musts contribute to the metabolic objective to balance their own cost. The optimality conditions give rise to general economic balance equations. Here, to describe the value structure by general laws, I wrote the optimality problems in an “expanded” form in which all dependencies between physical variables are described as explicit constraints. Economic values, dual to the physical variables, can then be defined by Lagrange multipliers. Just like their physical counterparts, economic variables are coupled between network elements (and via stoichiometric coefficients and elasticity coefficients, just like the physical variables). The value structure is described by simple economic rules, linking neighbouring economic variables. These rules are the basic laws of metabolic value theory. Their existence and structure can be proven for various kinds of models and model variables (e.g. growth rate or temperature) or objectives (e.g. fast cell growth in a whole-cell model), if the model can be written in expanded form. By combining the rules and assuming an optimal state^16^, we obtain balance equations that relate enzyme investments in networks to value production.

Metabolic value theory provides a new language for describing problems of function in cell biology, capturing both optimal and non-optimal states. It assumes a given metabolic objective, e.g. biomass production, and asks how each cell variable contributes to this objective. If an enzyme level contributes positively, there is an incentive for increasing this level, and we say that it has a positive use value. Since increasing the level is also costly, the enzyme also has a price (or “embodied value”). For each enzyme, embodied value and and use value must be balanced, so its total value must vanish. The same logic holds for all metabolic variables.

The value structure provides a new, local perspective on cell physiology. While metabolic optimality problems are formulated and solved in an abstract flux, enzyme, or metabolite space, the resulting economic variables can be visualised on the metabolic network itself (see Figure 5). To understand metabolism, we may therefore go in a circle: we start from network elements (such as enzymes and their reactants) and describe how they interact, e.g. how enzyme and reactant levels determine reaction rates and how these rates make metabolites accumulate. The network formed by these elements is the “space” on which metabolic dynamics take place and on which metabolic perturbations can be traced. To find optimal steady states, we need an even more abstract space, e.g. the space of possible flux distributions, and enzyme cost as a function on this space. This picture of cell states is less intuitive than our original network picture. But we need it because our optimality problems concern non-local, network-wide effects. Since all model variables are coupled, the benefit of an enzyme depends not only on its local neighbourhood, but on its effect on the overall metabolic state. However, surprisingly, these network-wide effects can again be written in a local form, namely as balances betwen neighbouring network elements. This brings us back to our network picture and closes the circle: metabolic value theory shows how network-wide optimality is locally reflected in single compounds and reactions.

**Figure 5.**
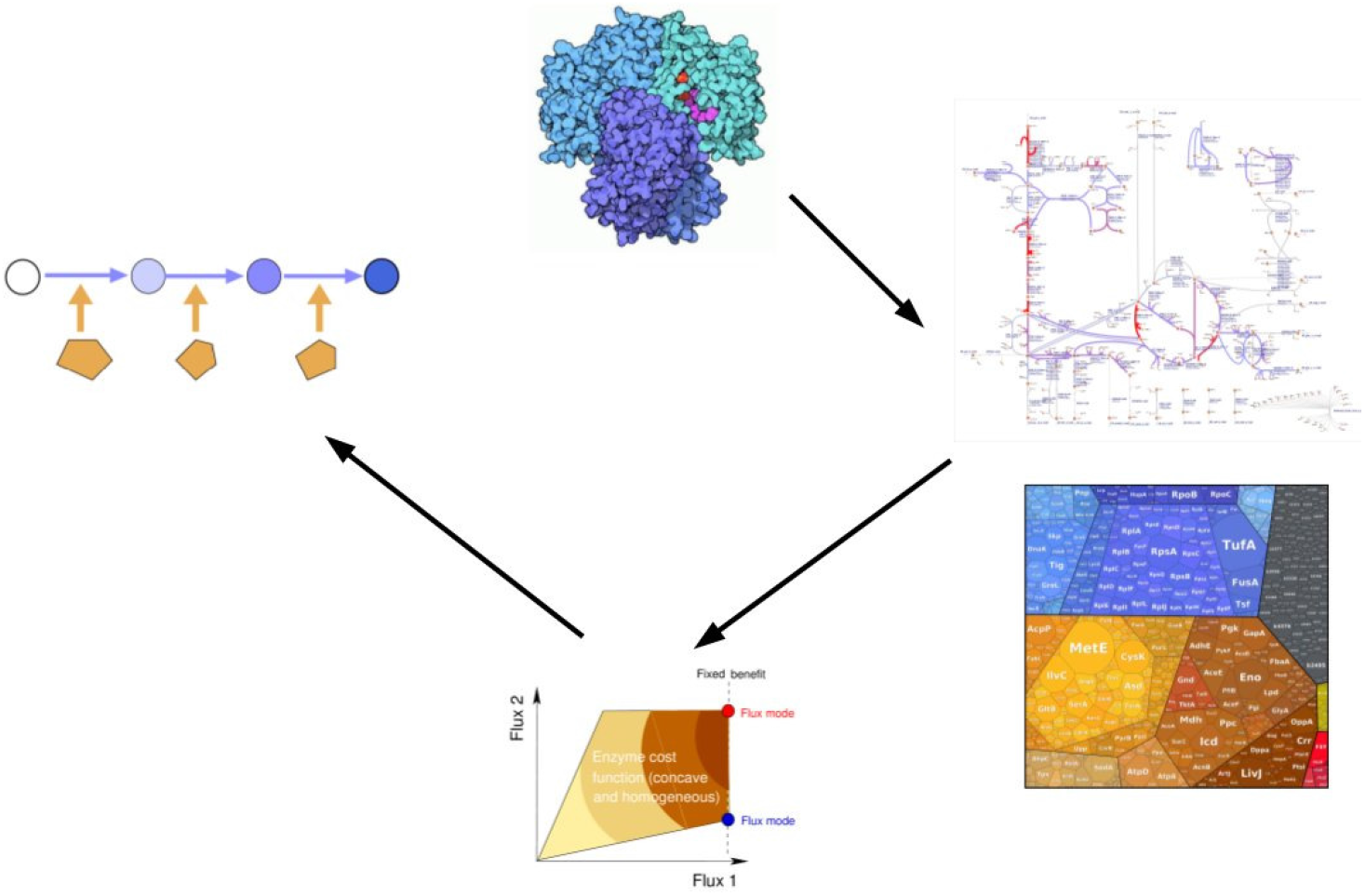
From biological networks to resource allocation problems and back. Molecular interactions (top) define metabolic networks. Fluxes and concentrations can be displayed on the network as “omics data”. To explain the shape of flux and enzyme profiles, we consider resource allocation problems. These problems are formulated in abstract spaces (e.g. flux space, bottom). Metabolic value theory (left) represents global fitness demands by the local economic values of metabolites, enzymes, and reactions. It brings us back to the metabolic network and shows how economic values depend on each other between neighbouring network elements.

The economic variables in Metabolic value theory can be compared between models. The theory also shows why certain features of optimal states (e.g. the absence of futile cycles or the balance between value production and enzyme investments) arise generally, for any choice of kinetic details and even in completely different modelling approaches (e.g. FBA or kinetic models), indicating that these variables and features are not model-specific, but exist in reality. How can such predictions be tested experimentally? We first need to defein what we mean by “fitness”, the function supposed to be maximised in optimal states. Generally, fitness refers to long-term survival (or a long-term selection advantage), but in experiments that select for fast growth (e.g. in long-term chemostats or serial dilution experiments), fitness can be understood to mean growth rate. To translate cell growth into cost and benefit functions for metabolic models, approximations are needed: for example, cell growth may be related to biomass/enzyme productivity by empirical formulae, or growth deficits due to enzyme demands may be estimated from known growth deficits measured for GFP overexpression. Alternatively, we may assume that cells maximise some target variable (e.g. ATP production) in the experimental set-up considered, and use this variable as a heuristic objective. Of course, our fitness function must correspond to a fitness defined operationally (in experiments). For example, with cell growth as a fitness function, protein costs may be quantified by assessing growth deficits caused by artificial protein expression [4, 8], or by measuring the effects of increased protein production costs (e.g. by nitrogen depletion or translation inhibitors). Next, we need experiments in which cells will behave optimally, even after a perturbation. Ideally, this would requires an evolution in the lab. Such experiments are not only required to explain actual adaptations, but also to measure economic variables such as enzyme prices or economic potentials. The key to defining economic variables lies in the choice of a fitness function, which then leads to a definition of marginal fitness values, i.e. fitness changes under variations of enzyme levels or import fluxes. With experiments to observe optimal states and state adaptations, we can quantify economic variables, check the validity of our economic laws, and validate their predictions. The idea is to compare different benefits and costs by “neutral replacements”. For example, to measure economic potentials, we may compare the import and biosynthesis of a metabolite and search for the break-even point, i.e. determine the possible transporter cost at which import and biosynthesis would cost the same. Assuming that our fitness is maximised in experiments, we can test some predictions of metabolic value theory. First, “high value” compounds (e.g. metabolites embodying large transporter or enzyme costs, or macromolecules embodying large machinery costs) should be used efficiently. Second, each enzyme investment should be balanced by a value production, given by multiplying the economic potential difference with the catalysed flux. Third, economic imbalances may lead to changes of enzyme levels during directed evolution, and may be reflected in rates of evolutionary change. To test this experimentally, one would have to measure precisely the enzyme costs in microbes, after an evolution in the lab [4, 23].

### 5.2 The value structure of metabolic states

Even if we accept an economic picture of cells, why does it have to be so complicated? Wouldn’t it be easier to count how much ATP is used in each cellular process and to define the cost of an enzyme or metabolite by the number of ATP molecules needed for its production? Can’t we express all costs and benefits simply in units of ATP? While such an “accounting” approach is certainly possible, it has strong limitations and does not answer many of the questions we can answer with MVT. First, cell processes are tightly linked and may have costly, indirect side-effects beyond the direct consumption of ATP; second, processes can act in circles (e.g. metabolite synthesis requires protein, and protein synthesis requires metabolites), so ATP consumption upstream or downstream of a given process may not be clearly distinguishable; and third, the cost of ATP itself (in terms of resource usage or growth deficit, or in comparison to other possible “currencies”) may be state-dependent. Finally, our main aim is not to describe *how* metabolic processes work, but *why* : i.e. what would be alternative, or possibly better strategies for the cell to work as a whole. Therefore, we cannot just describe how resources are spent in reality, but we need to ask how they *could be* spent, and whether this might lead to a higher fitness. To do this, we need a theory that does not only consider *absolute* resource balances, but balances of *marginal costs and benefits* (called “investment” and “value production”), describing the incentives for possible changes.

If metabolic optimality problems can be solved numerically, why do we need a general theory? Why should we treat Lagrange multipliers as “economic values” and why should we study their balance equations? There are several reasons. Numerical results are anecdotical: they only hold case by case and depend on specific parameter values; in contrast, the laws of metabolic economics hold generally and may explain metabolic behaviour even if kinetic details or parts of the network are unknown. Moreover, a general theory does not only predict *how* cells behave, but can also explain *why*. Finally, metabolic value theory gives metaphorical notions like “enzyme investment” a well-defined meaning and grounds them in realistic computational models. With its help, we can express beliefs about “economical behaviour of cells” more clearly and put them to a test.

### 5.3 From metabolic values to the economy of the cell

We learned that metabolism, besides its usual physical variables, has also a value structure, described by dual economic variables. The laws for these variables can be obtained from metabolic models with a fitness objective. We also saw that different models of the same system can lead to the same balance equations, which are therefore general laws. Metabolic Value Theory is not a new modelling framework, but a new perspective on existing frameworks which applies very broadly. While mechanistic models capture physical dynamics, economic values show us the “other side” – benefits and costs associated with this dynamics, described by dual variables. This other side can be studied for a wide range of models. In all cases, the economic variables reflect cost and benefit functions defined by the modeller. Metabolic value theory does not presuppose specific biological fitness objectives, but works for any objective functions, as long as cost, benefit, and bound functions are differentiable and if a local optimum exists. If optimality is assumed, we obtain local balance equations between cost and benefit; otherwise we obtain inequalities or equalities with “stress” terms that quantify mismatches between investment and value production.

Metabolic value theory can be applied to systems beyond metabolism, e.g. to models of entire cells. In this case, we treat all compounds (including lipids, mRNA, proteins, or DNA) as “metabolites” and describe their reactions (including macromolecule synthesis, modification, transport, and degradation) as parts of a larger network. Again, the value production principle must hold everywhere in this network: enzyme costs must be balanced by positive benefits, obtained through value production. For each mass-balanced compound, we can define an economic potential describing the compound’s contribution to cell fitness. However, in whole-cell models, the fitness looks different: if a cell model comprises protein production, no enzyme cost function is needed. Instead, costs for proteins and ribosomes arise automatically from energy and precursor consumption, from the need to produce these precursors, and from density constraints that limit the cellular capacity for these processes. Metabolic value theory allows us to cut out individual pathways, and use a local enzyme cost function that serves as a proxy for enzyme production, space demand, or missed opportunities in the surrounding cell.

## Abbreviations

FBA: Flux balance analysis,
LP: Linear programming,
MCT: Metabolic control theory,
MVT: Metabolic value theory.

## Acknowledgements

I thank Bernd Binder, Mariapaola Gritti, and Elad Noor for thinking with me about the topic. This work was funded by the German Research Foundation (Ll 1676/2-2).

## A Economic balance equations: short derivations

Economic balance equations can be derived in a few simple steps. Let us see this for the optimality problems shown in Figure 3. We write them all in expanded form: all model variables are described as free variables and all relations among them are written as explicit constraints. Furthermore, to make the problems comparable, we write them as *maximisation* problems. This means that lower bounds lead to positive Lagrange multipliers, resembling benefit terms (the variable should not be too low!), whereas upper bounds lead to negative Lagrange multiplier, resembling cost terms (the variable should not be too high! For more detailed derivations, see SI section **??**.

Before we come optimality conditions, let us think for a moment about constraints. Some constraints – for example, a required flux benefit – can be written in the form of bounds (i.e. inequality constraints) or in the form of fixed numerical values (i.e. equality constraints). If we consider a given optimal state, we have the choice: an active inequality constraint can always be replaced by an equality constraint, and an equality constraint can always be replaced by two inequality constraints. Sometimes, we can know in advance that an inequality constraint will be active (e.g. the flux benefit in flux cost minimisation): in the models below, I will formulate such constraints as inequalities to highlight their resemblance to cost or benefit terms (and the corresponding) sign of the Lagrange multiplier). In other cases (e.g. the kinetic rate law constraint)), we write equality constraints like we would write ≥ constraints) in the same order as a greater-equal constraint, with a constant number on the right.

### 1. Flux cost minimisation

In flux cost minimisation, we search for a stationary flux distribution **v** that minimises a flux cost function *a*(**v**) at a given flux benefit *b*(**v**) = *b′*. Optimality problem, Lagrange function, and optimality condition read:

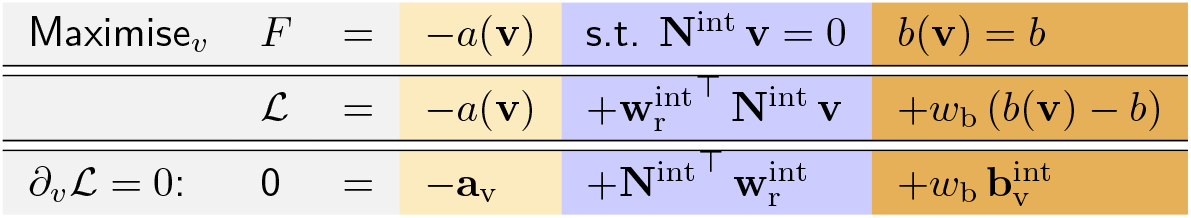

The optimality condition contains the flux price vector **a**_v_ = *∂a/∂***v**, Lagrange multipliers associated with stationarity constraints in a vector 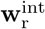, where 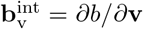 and *w*_b_ is the Lagrange multiplier for the flux benefit constraint. Colors are used to emphasise similarities with other optimality problems below. Setting 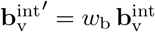 the resulting balance equation reads

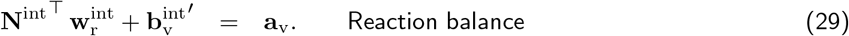

Note that this equation does not contain the original flux gain 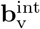, but the scaled version 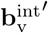, where the scaling (given by a Lagrange multiplier) varies between optimal states. If the flux benefit were not predefined, but included into the fitness function as a side objective (e.g. maximising *b*(**v**) − *a*(**v**)), we would obtain the same optimality condition, but this scaling factor would not appear.

### 2. Flux cost minimisation with enzymatic flux cost

Now we consider a kinetic model and minimise an enzyme cost *h*(**u**) at a fixed flux benefit *b*(**v**) = *b*; the fluxes must be stationary, the rate laws (between metabolite concentrations **c**, enzyme levels **u**, and fluxes **v**) must be respected, and the metabolite levels must satisfy the physiological bounds.

|  |  |  |  |  |  |  |
| --- | --- | --- | --- | --- | --- | --- |
| Maximise<sub>v,c,e</sub> | $F =$ | $-h(\mathbf{e})$ | s.t. $\mathbf{N}^{\text{int}} \mathbf{v} = 0$ | $\nu(\mathbf{c}, \mathbf{u}) = \mathbf{v}$ | $b(\mathbf{v}) = b$ | $\mathbf{c}_{\min} \leq \mathbf{c} \leq \mathbf{c}_{\max}$ |
| | $\mathcal{L} =$ | $-h(\mathbf{u})$ | $+\mathbf{w}_r^{\text{int}\top} \mathbf{N}^{\text{int}} \mathbf{v}$ | $\mathbf{a}_v^\top (\nu(\mathbf{c}, \mathbf{u}) - \mathbf{v})$ | $w_b (b(\mathbf{v}) - b)$ | $-\mathbf{g}_c^{\min\top} (\mathbf{c} - \mathbf{c}_{\min})$<br>$+\mathbf{g}_c^{\max\top} (\mathbf{c}_{\max} - \mathbf{c})$ |
| $\partial_v \mathcal{L} = 0:$ | $0 =$ | | $\mathbf{N}^{\text{int}\top} \mathbf{w}_r^{\text{int}}$ | $-\mathbf{a}_v$ | $+\mathbf{b}_v^{\text{int}}$ | |
| $\partial_c \mathcal{L} = 0:$ | $0 =$ | | | $+\mathbf{E}_c^\top \mathbf{a}_v$ | | $-\mathbf{g}_c^{\text{bnd}}$ |
| $\partial_e \mathcal{L} = 0:$ | $0 =$ | $-\mathbf{h}_e$ | | $+\mathbf{E}_e \mathbf{a}_v$ | | |

with Lagrange multipliers in vectors 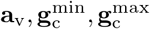 (with signs chosen for convenience). Compared to the previous problem, we obtain the additional Lagrange multiplier vectors **a**_v_ for rate law constraintsand 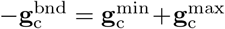 for metabolite bounds. Note that a metabolite can have a non-zero entry in 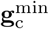 or 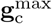, but not in both of them, and has vanishing entries for all inactive bounds. By solving the equations (in the order 3, 1, and 2), we obtain

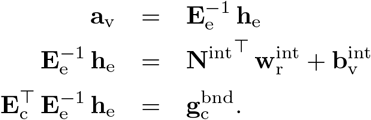

By combining the equations we obtain the economic balance equations

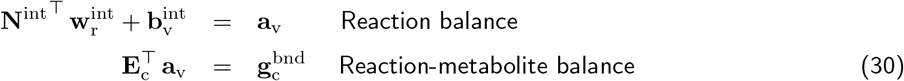

where the flux price vector **a**_v_, originally defined as a vector of Lagrange multipliers, is now simply given as 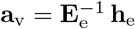.

### 3. Enzyme benefit-cost optimization

We consider a kinetic model and maximise the difference of a metabolic objective *b*(**v**) − *g*(**c**) and an enzyme cost *h*(**u**). All fluxes must be stationary, the rate laws must be respected, and the metabolite concentrations must the satisfy physiological bounds.

|  |  |  |  |  |  |
| --- | --- | --- | --- | --- | --- |
| Maximise<sub>v,c,e</sub> | $F =$ | $b(\mathbf{v}) - g(\mathbf{c}) - h(\mathbf{u})$ | s.t. $\mathbf{N}^{\text{int}} \mathbf{v} = 0$ | $\nu(\mathbf{c}, \mathbf{u}) = \mathbf{v}$ | $\mathbf{c}_{\min} \leq \mathbf{c} \leq \mathbf{c}_{\max}$ |
| | $\mathcal{L} =$ | $b(\mathbf{v}) - g(\mathbf{c}) - h(\mathbf{u})$ | $+\mathbf{w}_r^{\text{int}\top} \mathbf{N}^{\text{int}} \mathbf{v}$ | $\mathbf{a}_v^\top (\nu(\mathbf{c}, \mathbf{u}) - \mathbf{v})$ | $-\mathbf{g}_c^{\min\top} (\mathbf{c} - \mathbf{c}_{\min})$<br>$+\mathbf{g}_c^{\max\top} (\mathbf{c}_{\max} - \mathbf{c})$ |
| $\partial_v \mathcal{L} = 0:$ | $0 =$ | $\mathbf{b}_v^{\text{int}}$ | $+\mathbf{N}^{\text{int}\top} \mathbf{w}_r^{\text{int}}$ | $-\mathbf{a}_v$ | |
| $\partial_c \mathcal{L} = 0:$ | $0 =$ | $-\mathbf{g}_c$ | | $+\mathbf{E}_c^\top \mathbf{a}_v$ | $-\mathbf{g}_c^{\text{bnd}}$ |
| $\partial_e \mathcal{L} = 0:$ | $0 =$ | $-\mathbf{h}_e$ | | $+\mathbf{E}_e \mathbf{a}_v$ | |

By solving the equations (in the order 3, 1, and 2), we obtain

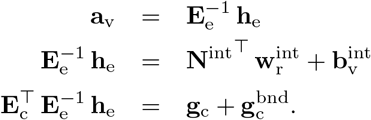

We obtain the same economic balance equations as before, but the terms have a different mathematical meaning:

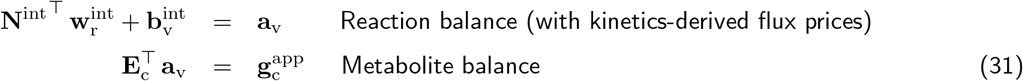

with the apparent metabolite price vector 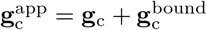 and again noting that 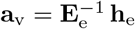.

### 4. Cost minimisation in metabolite space at given fluxes

We consider the same scenario as before, but now the fluxes are not optimised, but are predefined (and assumed to be thermodynamically feasible).

|  |  |  |  |  |
| --- | --- | --- | --- | --- |
| Maximise $c, e$ | $F =$ | $b(\mathbf{c}) - h(\mathbf{u})$ | s.t. $\nu(\mathbf{c}, \mathbf{u}) = \mathbf{v}$ | $\mathbf{c}_{\min} \leq \mathbf{c} \leq \mathbf{c}_{\max}$ |
| | $\mathcal{L} =$ | $b(\mathbf{v}) - h(\mathbf{u})$ | $\mathbf{a}_v^\top (\nu(\mathbf{c}, \mathbf{u}) - \mathbf{v})$ | $-\mathbf{g}_c^{\min \top} (\mathbf{c} - \mathbf{c}_{\min}) + \mathbf{g}_c^{\max \top} (\mathbf{c}_{\max} - \mathbf{c})$ |
| $\partial_c \mathcal{L} = 0:$ | $0 =$ | $-\mathbf{g}_c$ | $+\mathbf{E}_c^\top \mathbf{a}_v$ | $-\mathbf{g}_c^{\text{bnd}}$ |
| $\partial_e \mathcal{L} = 0:$ | $0 =$ | $-\mathbf{h}_e$ | $+\mathbf{E}_e \mathbf{a}_v$ | |

By solving the equations (in the order 2 and 1), we obtain

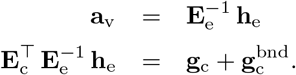

This yields the metabolite balance equation

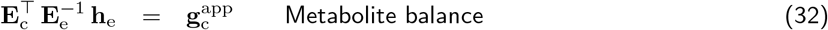

with the effective metabolite price vector 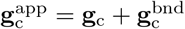.

### 5. Growth optimization

To model metabolism in a growing cell, we consider a kinetic model with enzyme levels as control variables and maximise the cell growth rate (a variable that affects dilution and enzyme cost). Fluxes must be stationary, rate laws must be respected, and metabolite concentrations must satisfy physiological bounds.

|  |  |  |  |  |  |
| --- | --- | --- | --- | --- | --- |
| Maximise $\lambda, v, c, e$ | $F =$ | $b(\lambda) - h(e, \lambda)$ | s.t. $\mathbf{N}^{\text{int}} \mathbf{v} - \lambda \mathbf{c} = 0$ | $\nu(\mathbf{c}, \mathbf{u}) = \mathbf{v}$ | $\mathbf{c}_{\min} \leq \mathbf{c} \leq \mathbf{c}_{\max}$ |
| | $\mathcal{L} =$ | $b(\lambda) - h(e, \lambda)$ | $\mathbf{w}_r^{\text{int} \top} (\mathbf{N}^{\text{int}} \mathbf{v} - \lambda \mathbf{c})$ | $\mathbf{a}_v^\top (\nu(\mathbf{c}, \mathbf{u}) - \mathbf{v})$ | $-\mathbf{g}_c^{\min \top} (\mathbf{c} - \mathbf{c}_{\min}) + \mathbf{g}_c^{\max \top} (\mathbf{c}_{\max} - \mathbf{c})$ |
| $\partial_\lambda \mathcal{L} = 0:$ | $0 =$ | $b_\lambda - h_\lambda$ | $-\mathbf{c} \mathbf{w}_r^{\text{int}}$ | | |
| $\partial_v \mathcal{L} = 0:$ | $0 =$ | | $\mathbf{N}^{\text{int} \top} \mathbf{w}_r^{\text{int}}$ | $-\mathbf{a}_v$ | |
| $\partial_c \mathcal{L} = 0:$ | $0 =$ | | $-\lambda \mathbf{w}_r^{\text{int}}$ | $+\mathbf{E}_c^\top \mathbf{a}_v$ | $-\mathbf{g}_c^{\text{bnd}}$ |
| $\partial_e \mathcal{L} = 0:$ | $0 =$ | $-\mathbf{h}_e$ | | $+\mathbf{E}_e \mathbf{a}_v$ | |

By solving the equations (in the order 4, 3, 2, and 1), we obtain

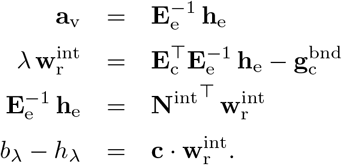

This yields the economic balance equations

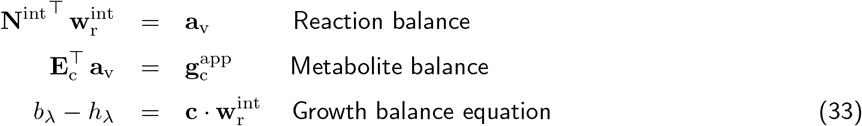

again with 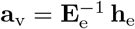. This time the apparent metablite price vector reads 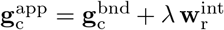, and there is no term 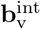. Note that third equation determines a scaling of all economic variables; if we are not interested in their absolute scaling, we can ignore this equation and do not have to know **c**. In the metabolite balance, the effective metabolite price reads 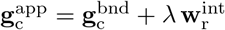.

### 6. Resource balance analysis (maximal growth)

A general linear RBA model can be obtained by slightly modifying the previous problem. Enzymes are not described as a separate type of compounds, but included in the compound vector **c**. As before, we maximise the growth rate, where fluxes must be stationary (“mass balance constraint”); linear capacity constraints must be satisfied (“capacity constraint”); weighted sums of the metabolite concentrations (e.g. in different cell compartments) must satisfy density constraints.

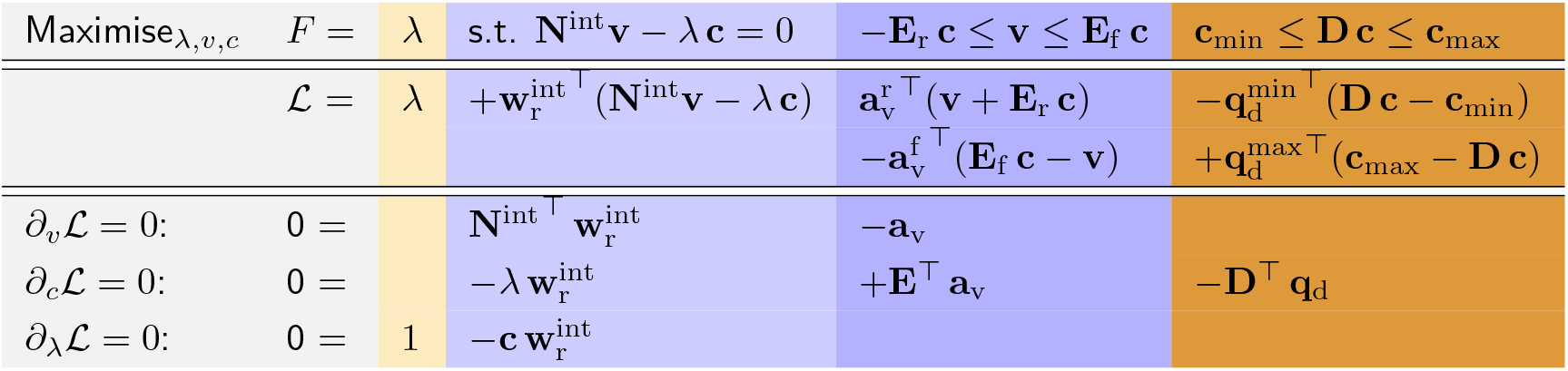

Here we defined 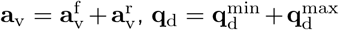, and a state-specific elasticity matrix **E** containing the elements of **E**^f^ for reactions that run in forward direction, and the elements of −**E**^r^ for reactions that run in backward reaction (and zero values for inactive reactions). Again there is no term 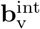. Since every active reaction will hit its capacity constraint [24], either 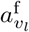 or 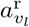 is non-zero for each reaction. This means that sign(**a**_v_) = sign(**v**) (with strict zeroes). By solving the equations (in the order 2,3,1), we obtain the economic balance equations

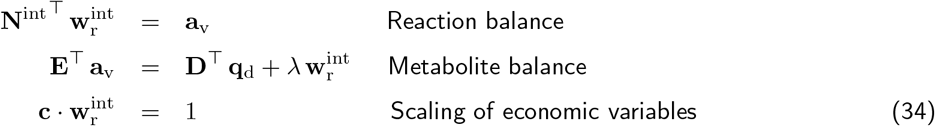

Given a solution of our optimality problem (the flux directions, active density constraints, metabolite concentrations, and growth rate), these equations can be used to determine the economic variables (see SI **??**).

### 7. Resource balance analysis (fixed growth)

In a variant of the RBA problem, we may describe a cell with a fixed (suboptimal) growth rate (e.g. in a chemostat), and which optimises some linear objective 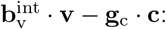:

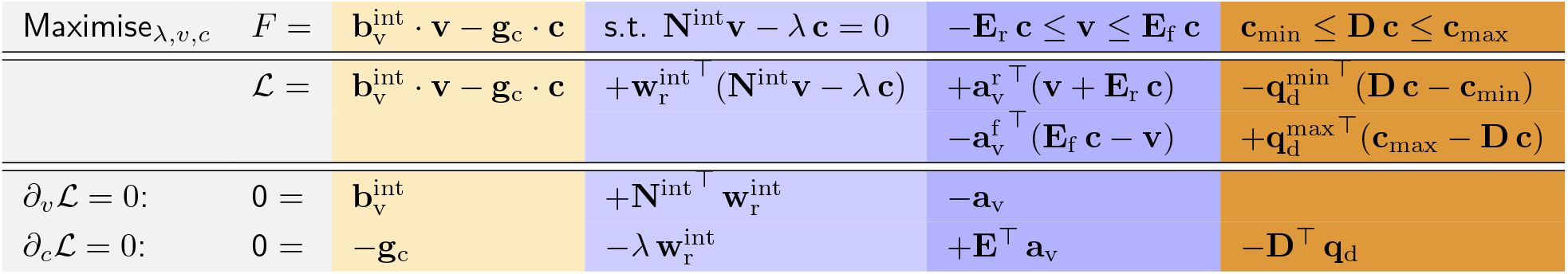

|  |  |  |  |  |  |
| --- | --- | --- | --- | --- | --- |
| Maximise$_{\lambda, v, c}$ | $F =$ | $\mathbf{b}_v^{\text{int}} \cdot \mathbf{v} - \mathbf{g}_c \cdot \mathbf{c}$ | s.t. $\mathbf{N}^{\text{int}} \mathbf{v} - \lambda \mathbf{c} = 0$ | $-\mathbf{E}_r \mathbf{c} \leq \mathbf{v} \leq \mathbf{E}_f \mathbf{c}$ | $\mathbf{c}_{\min} \leq \mathbf{D} \mathbf{c} \leq \mathbf{c}_{\max}$ |
| | $\mathcal{L} =$ | $\mathbf{b}_v^{\text{int}} \cdot \mathbf{v} - \mathbf{g}_c \cdot \mathbf{c}$ | $+\mathbf{w}_r^{\text{int}^\top} (\mathbf{N}^{\text{int}} \mathbf{v} - \lambda \mathbf{c})$ | $+\mathbf{a}_v^{\text{r}^\top} (\mathbf{v} + \mathbf{E}_r \mathbf{c})$<br>$-\mathbf{a}_v^{\text{f}^\top} (\mathbf{E}_f \mathbf{c} - \mathbf{v})$ | $-\mathbf{q}_d^{\min^\top} (\mathbf{D} \mathbf{c} - \mathbf{c}_{\min})$<br>$+\mathbf{q}_d^{\max^\top} (\mathbf{c}_{\max} - \mathbf{D} \mathbf{c})$ |
| $\partial_v \mathcal{L} = 0:$ | $0 =$ | $\mathbf{b}_v^{\text{int}}$ | $+\mathbf{N}^{\text{int}^\top} \mathbf{w}_r^{\text{int}}$ | $-\mathbf{a}_v$ | |
| $\partial_c \mathcal{L} = 0:$ | $0 =$ | $-\mathbf{g}_c$ | $-\lambda \mathbf{w}_r^{\text{int}}$ | $+\mathbf{E}^\top \mathbf{a}_v$ | $-\mathbf{D}^\top \mathbf{q}_d$ |

where 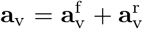 and 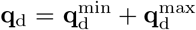 as before. Again, by solving the equations (in the order 2,3,1), we obtain the economic balance equations

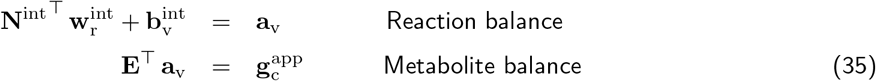

where, this time, 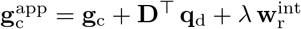.

All the problems shown lead to very similar balance equations. However, these are only a few examples. By replacing constraints by optimality criteria (or the other way around), or by considering multi-criteria optimisation, we can construct many more variants of the optimality problems, but the balance equations remain the same.

## B. Optimality problems on networks

To obtain a theory of optimal metabolic states, based on variations and shadow values, we consider optimality problems on networks more generally. The variations on our network – in our specific case, in the variables (**v, c, e**) – are described by a vector **x**.The feasible vectors **x**, satisfying the model constraints, form an algebraic manifold, and economic values follow from cost and benefit gradients on this manifold. An objective *f* defines direct values (by its gradient in state space), total values (by its gradient in the tangential space of the manifold, comprising the total derivatives), and indirect values (by the difference of the two). To see this in detail, we consider a general network model in an expanded form. The variables *x*_*i*_ are constrained by dependencies *x*_*i*_ = *g*_*i*_(**x**), which must be acyclic: a variable *x*_*i*_ can depend only on variables *x*_*j*_ with indices *j < i* (and variable *x*_*i*_ is called a “child” of *x*_*j*_.). If dependencies are depicted by arrows, these arrows form a directed acyclic graph (DAG). Aside from these dependencies, variables can also be constrained by bounds and fixed values (as equality and inequality constraints) and are scored by a fitness function *f* (**x**). Our model in general form reads

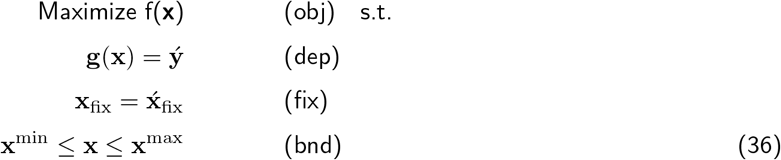

In the following, we consider optimal states and simplify the optimality problems in two ways: first, any general inequality constraints *g*(**x**) ≤ 0 are replaced by an assignment *g* = *g*(**x**) and an inequality *g* = 0. Second, all inequality constraints are either omitted or replaced by equality constraints, without chaning the optimal state.. If a constraint can be omitted without changing the optimal state, the constraint is called “ineffective” In general, in a given optimal state, inactive inequality constraints will have no effect, and active inequality constraints can be replaced by equality constraints. Therefore, in this state, we can ignore all inactive bounds and replace all active bounds by fixed values. Therefore, for simplicity, we now limit ourselves for problems with equality constraints only. Geometrically, we can see the feasible states **x** as points on a state manifold, and the fitness *f* as a function on this manifold [13]. If our point is varied by an “infinitesimal” *δ***x** (in fact, a variation described by a differential form), the fitness variation is given by by *δf* = **f**_**x**_ *δ***x**, where **f**_**x**_ is the gradient of the fitness function. If a state manifold is differentiable (which holds for our models [13]), variations within the manifold can be described as variations *δ***x**_∥_ in the tangential space. Mathematically, infinitesimal variations around a reference point (the state of interest) can be described by differential forms *δ***x**, and their fitness effects are given by the scalar product *δf* = **f**_**x**_ *· δ***x** with the help of the fitness gradient **f**_**x**_ in the reference point **x**. Intuitively, we can see fitness gradients as forces pulling towards regions of higher fitness. Mathematically, “pulling” means that there exist feasible variations *δ*_∥_**x** (within the tangential space) that woudl increase fitness, i.e. variations for which **f**_**x**_ *· δ*_∥_**x** is positive. In each point of the manifold, the fitness gradient **f**_**x**_ consists of a tangential component 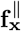 (within the tangential space) and an orthogonal component 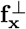 (in the orthogonal space). The first component pulls the state towards higher fitness values *within the manifold*, while the second component pulls in a direction that is prohibited by the constraints. In the optimal point, the fitness gradient **f**_**x**_ is orthogonal to the state manifold and its tangential component 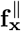 vanishes. There is no variation that would increase fitness; or in other words, in the optimal point the state manifold and the fitness gradient are orthogonal to each other. In terms of our original problem, the gradient **f**_**x**_ describes direct values, the tangential vector 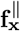 the total values, and the negative orthogonal vector 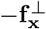 (i.e. total value minus direct value) the indirect values.

Let us now come back to the full problem with inequality constraints. Solutions of the constrained optimality problem Eq. (36) must satisfy the Karush-Kuhn-Tucker conditions. We define the Lagrangian

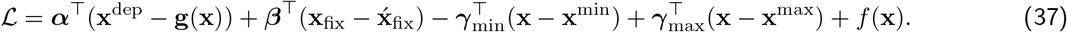

With the vector **f**_x_ (“direct economic values”) and matrix **Γ** = (*∂***g***/∂***x**)^*T*^ (“connection matrix”), the optimality condition reads ∇_**x**_ ℒ = 0, i.e.

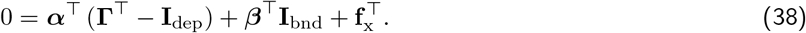

The matrices **I**_dep_ and **I**_bnd_ are obtained from an identity matrix **I** (for internal metabolites) by selecting all rows that correspond to dependent variables or variables with active bounds, respectively. Transposing this equation yields

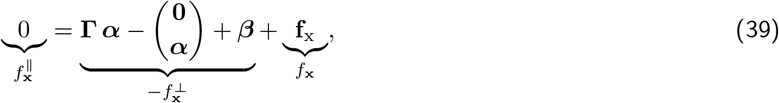

with subvectors **0** and ***α*** in the second term (a column vector) for control variables and derived variables, respectively. As shown above, the terms in (39) can be associated with the fitness gradient *f*_**x**_, its tangential component 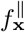, and its perpendicular component 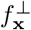 (see Fig. **??**). Separating the rows for choice variables and derived variables, we obtain

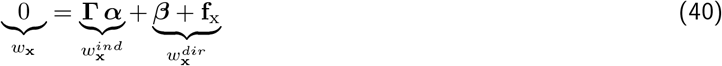

for choice variables (such as **e** and **c** in our standard kinetic models) and

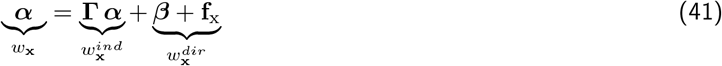

for dependent variables (such as **v** and **r** in our standard kinetic models). These are our balance equations (with terms derived from fitness derivatives and Lagrange multipliers). By renaming our variables we obtain a the “economic law”

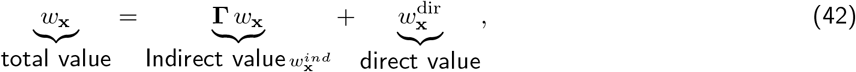

where the total value *w*_**x**_ of each control variable must vanish (see Eq (39))! Written in this form, terms are named by their mathematical origin, but only by their roles in the formula (total value, direct value, or indirect value acquired from the values of their child variables). These are the economic rules in their general form. There are two other details. First, a “direct value” can comprise a direct derivative and a shadow value, (where the shadow values of inactive bounds are zero. Second, non-optimal states can be described in almost the same way. In optimal states, the tangential gradient 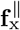 vanishes, but in general non-optimal states, there remains a tangential gradient 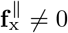 (called stress **t**_x_). Moving the stress term to the left, we obtain the general economic rule (for non-optimal states and with explicit shadow values from bounds)

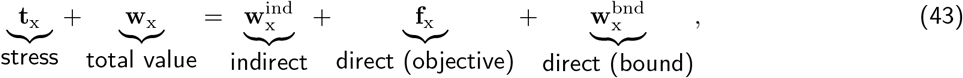

with an indirect value (or “load”) 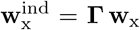 and a direct value (or “gain”) 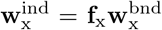. Of course, our previous equations can be reobtained by omitting some terms. For example, setting the stress **t**_x_ to 0 yields the balance equation for an optimal state; by setting **w**_x_ = 0, we obtain the economic rule for a choice variables (e.g. enzyme levels in the standard kinetic model); and if a physical variable has no direct value, then the associated economic value, in optimal states, is given by

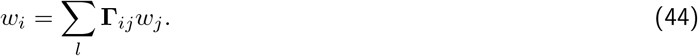

In this case, the economic value (of *x*_*i*_) is entirely acquired from the values of its child variables *x*_*j*_.

## Footnotes

1 In our metabolic “standard models”, the vector **e** contains enzyme concentrations. However, in other models it may also contain other control variables, such as mRNA levels, that appear as prefactors in rate laws.

2 Derivation: 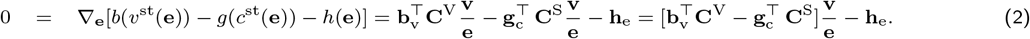 where **v***/***e** stands for the enzyme elasticity matrix **E**_e_ = Dg(**v**)Dg(**e**)^−1^.

3 If *x* is (directly) constrained by bounds *x*^min^ ≤ *x* ≤ *x*^max^ or by a predefined value *x* = *x*^fix^, this leads to a shadow price, which plays a similar role as *f*_*x*_ and can be added to it.

4 In a linear pathway with thermodynamic forces *θ*_*l*_ and *K*_M_ values set to 1, the substrate elasticities are larger than the product elasticities [14]. According to the balance equation, around a metabolite, the enzyme investment in the producing reaction should thus be larger than in the consuming reaction. Since this holds for each metabolite, the enzyme investments must decrease along the chain.

5 A nonlinear benefit function with gradient *b*_**v**_ = *∂b/∂***v** would lead similar optimality conditions. Here we consider linear functions just for simplicity.

6 In models with moiety conservation (e.g. if ATP and ADP appear in reactions only as cofactor pairs, and therefore [ATP]+[ADP]=const.), the concentrations of conserved moieties can be treated as model parameters.

7 In dynamically unstable states or bifurcation points, our theory does not apply. In a metabolic system, the enzyme profiles will not always uniquely determine the metabolic state. On the one hand, the state may also depend on conserved moiety concentrations (which follow from the initial conditions). On the other hand, in cases of multistability, we need to select one of the possible steady states.

8 For simplicity we assume bounds on individual metabolites. Density constraints for a sum of metabolite concentrations can be treated similarly and will be discussed below.

9 In a dynamical reaction system, dilution alone would lead to an exponential concentration decrease. Sine-wave oscillations of enzyme and metabolite concentrations can be described by complex-valued exponential functions. This resembles an example in physics: an overdamped pendulum shows an exponentially decreasing motion, which is mathematically related to the motion of oscillating pendulum because sine and cosine functions are exponential functions with an imaginary function argument.

10 Here we consider deterministic, non-spatial network models in steady state. Other possible variables (which can be included in the theory) are for instance compartment sizes, electric potentials, or temperature.

11 In some models such terms play a role. External concentration prices may matter in modular models, in which models are coupled (with external metabolites between them) and an optimal compromise (for the external metabolite concentrations) needs to be found. For example, one submodel produces ATP and the other submodel consumes it; the first submodel would run more efficiently at a lower ATP concentration, while the second submodel would run more efficiently at a higher ATP concentration. In the optimal state, the (positive) ATP price from the first model and the (negative) ATP price from the second model must exactly cancel out.Similarly, internal production gains matter in models with dilution or could be needed to penalise loss by diffusion (in models in which diffusion reactions are not explicitly described).

12 Dual variables also play a role in physics. In thermodynamics, the chemical potentialsare defined as derivatives of the Gibbs free energy with respect to substance amounts (in moles), and are dual variables of the substance amounts (the physical variables). According to the second law of thermodynamics, thermodynamic systems tend towards states of minimal Gibbs free energy. The chemical potentials are a useful concept to describe this. The chemical potential *µ*_*i*_ measures the Gibbs free energy contained in one mole of substance *i*. Accordingly, a change in the mole number *δn*_*l*_ leads to an energy change *δG* = ***µ*** *·* **n**. Knowing the chemical potentials, we also know which processes would dissipate Gibbs free energy (by changing the substances’ mole numbers), and may therefore happen spontaneously.

13 Whenever a variable hits a bound, the corresponding shadow value acts as a gain or price, creating an extra term in the value balance, and therefore an additional value “inflow” or “outflow”.

14 Formally, like in the case of variables hitting a bound, this term can also be seen as virtual value inflow or outflow; together with this term, value conservation would hold again at least formally.

15 Interestingly, variational principles in physics used are often not maximisation principles but extremality principles, i.e. they may be satisfied by maximal, minimal, or saddle point solutions. In contrast, if we apply variational principles to living systems, we typically know whether a target should be minimised or maximised. We assume that cells maximise, for example, a benefit or minimise a cost on a set of possible cell states. Optimising a simple objective can be seen as an approximation of more complicated multi-objective problems, which would be a more realistic description of cells.

16 Metabolic value Theory also applies to non-optimal states. In optimal states, investments and value production are balanced. However, real cells are never exactly optimal. If an objective function has been defined, deviations from optimality can be expressed by “stress” terms in the cost-benefit balance equations.

